# Multimodal Data Fusion Reveals Morpho-Genetic Variations in Human Cortical Neurons Associated with Tumor Infiltration

**DOI:** 10.64898/2025.12.26.696632

**Authors:** Yufeng Liu, Zhixi Yun, Lingli Zhang, Wen Ye, Kaifeng Chen, Xiefeng Wang, Mengzhu Ou, Jing Rong, Xiaomin Yang, Lei Mao, Liang Chen, Nan Ji, Yongping You, Chiyuan Ma, Ying Mao, Junxia Zhang, Liwei Zhang, Hanchuan Peng

## Abstract

We introduce **LetsACT** (Light-Electron-Transcriptome synergistic ACTomography), a multimodal integration platform that overcomes the limitations of single-modality data acquisition and analysis of human brain cells while synergistically leveraging the strengths of each modality. Our approach enables the rapid sample preparation, cell injection, imaging, and multimodal integration of large-scale human neuronal datasets at single-cell resolution. By generating initial laser-scanning-microscopy based optical reconstruction of neuron morphologies followed by refining them using electron-microscopy derived morphological priors, we have assembled one of the largest human cortical morphology datasets to date: 8,398 neurons from 58 donors, with high cortical coverage. This platform is then applied to studying morphological impact of tumor infiltration. Pyramidal neurons in glioma-infiltrated tissues display clear volume shrinkage in somas and branches, tapering from the soma to nearby dendritic compartments. By integrating these morphological variations with spatial and bulk transcriptomic profiles, we find that glioblastoma tissues exhibit dysregulation of 15.29% of genes, including overexpression of TERT, whereas infiltrated tissues show 7.74% gene dysregulation, characterized by overexpression of tumor suppressors such as CDKN2A and TP53. Our analysis implies that pyramidal neurons observed in these infiltrated tissues may involve an active defense instead of undergoing passive apoptosis. Our finding also indicates that **LetsACT** establishes a valuable resource for the large-scale, comprehensive morpho-genetic analysis of human tissues.

## Introduction

While genetic signatures that distinguish human tumor and tumor infiltrated tissues from normal tissue have been extensively characterized (e.g, Aran et al., 2017), the integration of morphological and genetic features, particularly in the human brain, has remained elusive because of technical challenges. In particular, there has been a lack of scalable morphological profiling approaches capable of capturing complex three dimensional neuronal morphologies in human brain tissue at single cell resolution. Although post mortem analyses have yielded valuable baseline information, examinations of fresh surgical tissue preserve native cellular morphology and molecular signatures that may be degraded in post mortem samples, thereby providing real time snapshots of tumor and brain interactions. Importantly, this approach also has direct clinical relevance: the combined analysis of morphology and genetics in fresh surgical specimens can inform personalized treatment strategies and offer prognostic insights that post mortem studies cannot provide.

Morphological screening of human brain tissue presents substantial technical and biological challenges. Unlike model organisms, where *in vivo* genetic labeling and imaging are routine (Ghosh et al., 2011; Kuramoto et al., 2009; Lin et al., 2018; Luo et al., 2016), human brain tissue must be processed outside the body while maintaining sufficient cellular viability for labeling. Access to human tissue is limited and unpredictable, which requires protocols that are rapid and robust and that can accommodate variable time intervals after surgery, diverse pathological states, and heterogeneity between individuals. The thick and lipid rich nature of human brain tissue also impedes both dye penetration and optical clarity, and the absence of standardized morphological atlases for human neurons further complicates cell type identification and quality control. Together, these factors have produced a major gap in our understanding of human neuronal morphology, especially in disease contexts in which structural alterations may accompany genetic changes.

Achieving statistically robust morphological characterization of human neurons requires an unprecedented scale in sample size as well as regional and demographic coverage, and this is difficult to achieve in human studies. Unlike transcriptomic analyses, where thousands of cells can be profiled from a single sample, morphological reconstruction requires many hours of manual or semi-automated single cell tracing in order to capture complex arborization under sparse labeling conditions. Truly comprehensive sampling must include multiple cortical regions and multiple neuronal classes, since morphology varies widely from densely branching nonpyramidal interneurons to highly elaborate pyramidal cells, while also ensuring balanced representation across ages and sexes to account for age related changes and sex specific differences. Additional logistical challenges include coordinating tissue collection across multiple centers, maintaining consistent protocols despite variable surgical schedules, and achieving demographic balance within the constraints of neurosurgical practice. Previous studies, which have typically been limited to hundreds of neurons from only a few donors (Berg et al., 2021; Mohan et al., 2023), face difficulty in distinguishing true biological variation from technical noise.

Large-scale reconstruction and screening of human neurons need to overcome several fundamental technical challenges. While recent studies have advanced human neuron techniques and resources (Ataman et al., 2016; Buchin et al., 2022; Jacobs, 2001; Loomba et al., 2022; O’Leary et al., 2021; Planert et al., 2025; Scholtens et al., 2022; Shapson-Coe et al., 2024; Zaslavsky et al., 2019), Three-dimensional (3-D) reconstruction of the complicated arbors of neurons remains limited by labor-intensive manual annotation. Automatic neuron tracing primarily developed for model organisms shows limited robustness for human neurons (Feng et al., 2015; Li et al., 2019; Peng et al., 2011; Quan et al., 2016; Xiao & Peng, 2013). Indeed, human neuronal images may present unique challenges, including elevated noise, reduced signal in distal branches (**Fig. S1**), and elongated thin tubular structures that current segmentation algorithms struggle to capture accurately (Kirchhoff et al., 2024).

In this study, we developed Light-Electron-Transcriptome synergistic ACTomography (**LetsACT**), a high throughput platform that integrates light microscopy (LM), electron microscopy (EM), and transcriptomics for large-scale reconstruction and analysis of human neurons. We enhanced ACTomography (Han et al., 2023) with an automated multi-needle injection system (Peng et al., unpublished), enabling sparse and high-throughput labeling of human neurons through somatic dye injection guided by bright-field or differential interference contrast (DIC) microscopy (**Fig. S1a**). To address the inherently noisy and fragile nature of human imaging data, we established an automated reconstruction pipeline with adaptive enhancement algorithms that selectively strengthen distal branches, reduce proximal noise, and improve continuity of neurite segments. Primary topology was further refined using spatial distribution priors derived from EM based reconstructions. LetsACT enabled construction of ACT-H8K, a large dataset of 8,398 reconstructed human neurons from 58 donors (32 male and 26 female) spanning diverse age groups and brain regions. The resulting resource represents both a quantitative expansion and a qualitative advance in the definition of human neuronal morphology. Beyond morphological characterization, our findings reveal divergent responses of neuronal subtypes under oncogenic stress. The morphology of pyramidal neurons shows a non-uniform, distance dependent attenuation from the soma to nearby dendritic compartments. By integrating structural measurements with multi-scale transcriptomic profiles across normal and glioma infiltrated cohorts, we provide evidence that these morphological shifts may represent an active defensive strategy rather than passive apoptotic collapse. In summary, this work establishes LetsACT as an important platform for linking cellular anatomy with molecular pathology in the human brain.

## Results

### Assemble dendritic morphologies of 8,398 human cortical neurons

We collected 65 fresh surgical samples and one postmortem sample from 58 donors (**Supplementary Table S1**). These tissues were sectioned into 697 slices, each of which has a thickness between 200 μm to 600 μm. These sections are thicker than that of H01 (170 μm, Shapson-Coe et al., 2024), which is another large, published dataset of human neurons’ 3-D morphology. In our LetsACT approach, individual neurons were labeled via ACTomography (Han et al., 2023) by microinjecting visible somas with Lucifer Yellow (LY). 3D image stacks of neurons were then acquired using two-photon laser scanning microscopy (**Fig. 1a–b**; **Fig. S1a**) (**Methods**). 8,398 neurons’ 3-D dendritic morphologies, referred to as ACT-H8K, were digitally reconstructed and evaluated (**Fig. 1c–e**) using an AI-powered approach explained below.

**Fig. 1.**
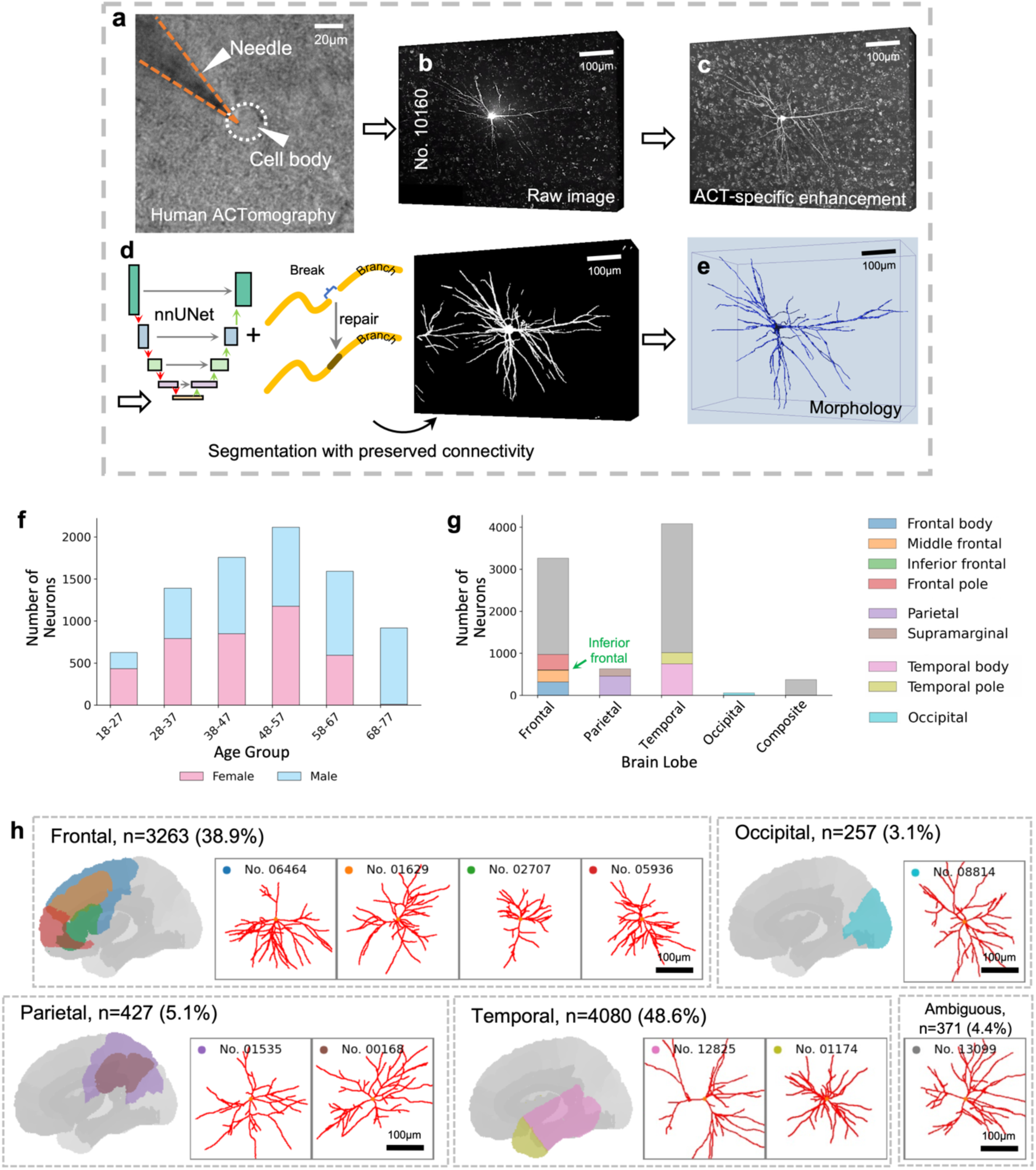
Automated tracing of 8,398 human cortical neurons. **a-e,** The automatic initial tracing pipeline in LetsACT. **f,** Age and gender distribution of the reconstructed neurons. **g,** Neuron composition by brain regions. Composite: neurons derived from tissues spanning two cortical lobes. The gray bars in each lobe indicate brain areas situated along the boundary between two adjacent regions. **h,** The reconstructed neuron morphologies. Left, brain regions are highlighted on the Yale Brain Atlas. Right, exemplar morphologies from each of these regions. The neuron IDs are displayed at the top. The colors of the top-left dots indicate the brain regions from which the neurons originate.

ACT-H8K covers a broad range of ages and genders (**Fig. 1f**), and all four cortical lobes. In this study we focused on 9 sub-lobar regions out of all 20 cortical regions defined in the Yale Brain Atlas (McGrath et al., 2022) (**Fig. 1g**). These neurons exhibit arborizations across different regions (**Fig. 1h**), with the majority originating from the temporal lobe (48.6%) and frontal lobe (38.9%). The average age of subjects was 49 years, with ages spanning 18 to 75 years, and the gender distribution was nearly balanced, with 32 males and 26 females (**Supplementary Table S2**).

We manually annotated neurons into three categories: pyramidal (excitatory), nonpyramidal (mostly inhibitory), and uncertain, based on cellular image and morphology (**Supplementary Table S3**). Each neuron was evaluated by 2 or 3 annotators. Only those annotations achieving unanimous independent consensus were retained (pyramidal and nonpyramidal). Cortical layer identity could be inferred from DAPI-based cytoarchitecture or morphological properties of somas and dendritic arborizations (Yun et al., 2024). Notably, glioma cells are reported to disrupt both spatial organization and neuronal circuits (Civita et al., 2020; Krishna et al., 2023), which makes laminar identification in tumor or tumor-infiltrated tissues inherently less reliable, and thus not a focus in this study.

To tackle weak or uneven signals in the presence of strong background noise such as the haze often seen in the soma-region and autofluorescence possibly associated with blood (**Fig. S1**), we built an automated neuron reconstruction pipeline with three stages: image enhancement, segmentation, and topology-preserving tracing (**Methods**). Essentially, for enhancement, we modeled the haze around each soma as a Gaussian diffusion field and applied a spatially differentiated gamma correction (ACT-specific enhancement, **Fig. 1c**; **Fig. S2**), which suppresses proximal artifacts while amplifying distal neurites. For segmentation, we trained an artificial neural network model with a connectivity-enhancing loss to improve neurite continuity (**Fig. 1d**). During tracing, somas were detected and handled separately to enforce first-order connectivity between primary stems and somas. We refined the first-order dendritic topologies by incorporating priors shown subsequently. Branch radii were estimated by combining reconstructed skeletons with local intensity profiles. All the reconstructions were standardized to the isotropic coordinate space and are publicly available for the community as one of the largest human neuron datasets with extensive anatomical and morphological diversity (**Data availability**).

### Refine 3D neuron morphology by integrating electron-microscopy priors

Morphological priors from “gold-standard” datasets are critical for neuron reconstruction, especially for morphologies derived from high-resolution electron microscopy. We observed that initial reconstructions of ACT-H8K (ACT-H8K-I) may suffer from artifacts, such as excessive primary stems (red arrow in **Fig. 2b**). In LetsACT, we incorporated electron-microscopy priors into our light-microscopy data by first merging the first-order branches (primary stems) and then evaluating overall similarity. We compared the distribution of primary stem numbers across four datasets of neuronal dendrites generated using distinct methodologies: the ACT-H8K-I dataset (*n*=8,398), pyramidal neurons from the middle temporal gyrus (MTG) in the DeKock v1.1 dataset (*n*=89; Mohan et al., 2023), the Allen human cell types dataset (*n*=297; Berg et al., 2021), and morphologies from the electron microscopy dataset H01 (EM data from MTG; *n*=3,446). The original H01 dataset (C3 segmentation layer) contained skeletons of 12,517 neurons, most of which were fragmented. We developed a large-language-model (LLM) based method (Gu et al., unpublished) to refine this dataset, yielding 1,598 high-confidence neurons (*level 1*) and 1,848 moderate-confidence neurons (*level 2*), resulting in a total of 3,446 neurons used in this study. Our findings revealed that neurons in the DeKock, Allen, and H01 datasets consistently exhibited no more than 12 primary stems per neuron. In contrast, ACT-H8K-I reconstructions demonstrated a considerable percentage of polystemic reconstructions, with 2.9% of neurons (*n*=243) possessing more than 12 primary stems (red arrow in **Fig. 2b**).

**Fig. 2.**
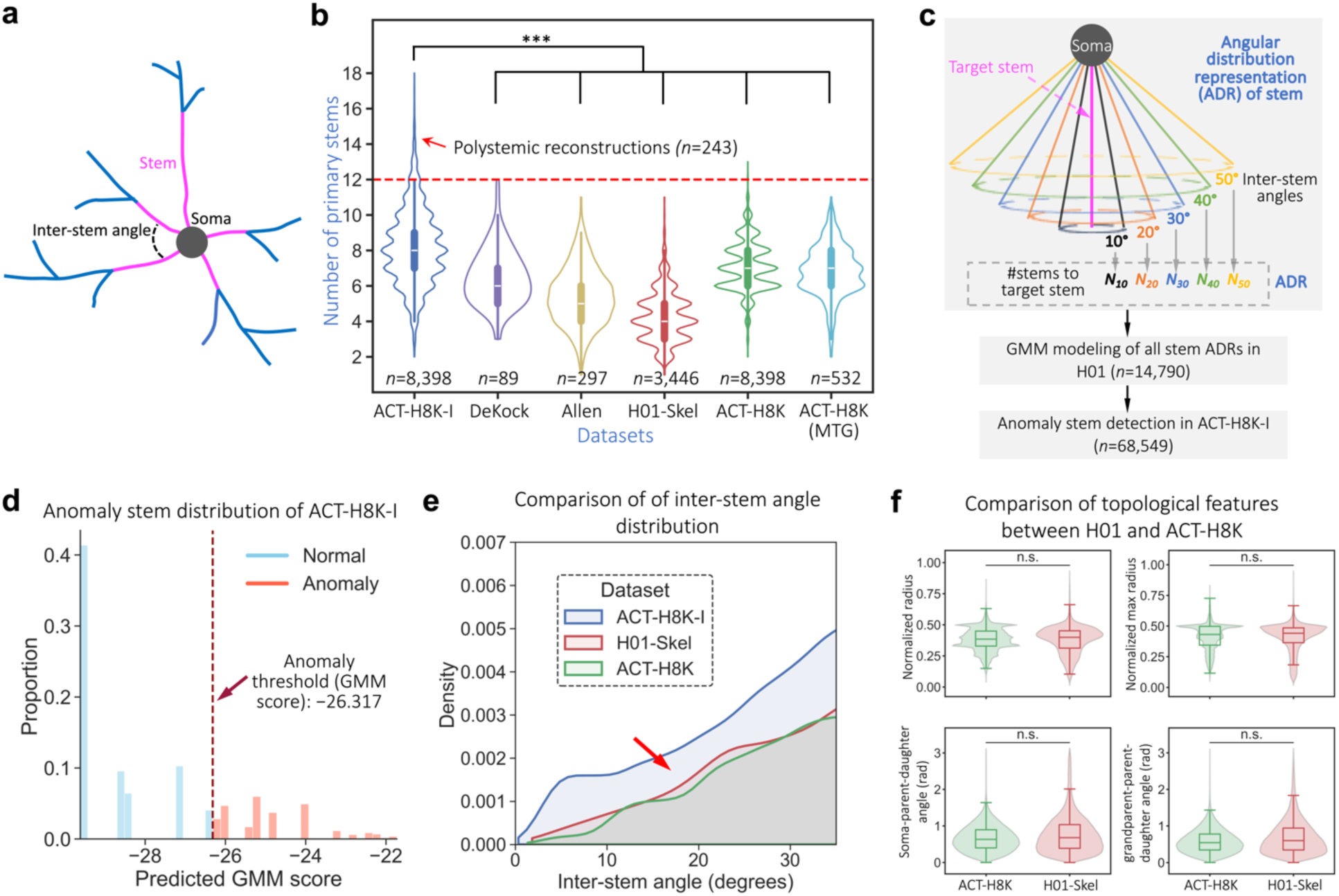
EM-LM synergized refinement of neuron structures. **a**, Schematic of a neuronal dendrite, with highlight of primary stem and inter-stem angle. **b**, Distribution of dendritic primary stem numbers across multiple datasets (ACT-H8K-I, DeKock, Allen, H01-Skel, ACT-H8K, and MTG neurons in ACT-H8K). The number of polystemic reconstructions (*n*=243) that contain over 12 stems are annotated. Statistical significances are estimated using two-sided Mann-Whitney U tests: \*\*\**p*-value < 0.001. Violin plot: edges, the kernel density estimation of the data distribution; central line, the median; inner box, the interquartile range (25th to 75th percentiles). **c**, Schematic illustrating the angular distribution representation of a primary stem, along with the process of anomaly detection using a GMM trained on stems in H01-Skel. **d**, Stem anomaly distribution of the initial ACT-H8K reconstructions. 28.1% of stems from ACT-H8K-I reconstructions were identified as anomalies, using a threshold (-26.317) calculated as the 5th percentile of the ADR distribution in the EM dataset. **e**, Density plot of inter-stem angle distributions in ACT-H8K, compared with H01-Skel, and ACT-H8K. The red arrow highlights the reduced density of small inter-stem angles. **f**, Comparison of topological features between ACT-H8K neurons and H01 neurons. “n.s.” indicates non-significance, defined as a two-sided Mann-Whitney U test *p*-value > 0.05, or an effect size in range [0.45, 0.55]. The feature “normalized radius” denotes the ratio of the radius of the current branch to the sum of the radii of the current branch and its parent branch. “Normalized max radius” is similar but uses the ratio of the maximum radius among all nodes in the branch. “Soma-parent-daughter angle” denotes the angle between the vector from the soma to the initial node of the branch and the vector from the initial node of the branch to its terminal node. “Grandparent-parent-daughter angle” is the angle between the vectors of two successive branches. Box plot: edges, 25th and 75th percentiles; central line, the median; whiskers, 1.5×the interquartile range of the edges; dots, outliers.

We developed an EM-LM synergized refinement for primary stems, utilizing Gaussian Mixture Model (GMM) analysis to model the spatial distribution of primary stems in the H01 dataset. Specifically, for each primary dendritic stem in the EM dataset (*n* = 14,790 stems), we quantified the number of neighboring stems falling within angular bins of 10°, 20°, 30°, 40°, and 50°, thereby constructing a 5-dimensional angular-distribution representation (ADR; **Fig. 2c**). To prioritize small-angle artifacts, the weights assigned to these five bins were progressively decayed by a factor of 0.1. A GMM with 11 components (**Fig. S3a**) was trained on these ADRs and subsequently applied to the primary stems of ACT-H8K-I (*n*=68,549). The model classified 28.1% of stems in ACT-H8K-I as anomalies (**Fig. 2d**). We also observed that the proportion of anomalous stems correlated with the total number of stems per neuron, with neurons containing seven stems exhibiting low level of anomaly rates (**Fig. S3b**).

To rectify, we implemented an iterative process to correct anomaly stems by merging them with the spatially closest stems (**Methods**), yielding the ACT-H8K dataset. This detection-and-merging procedure continued until either the stem count per neuron was reduced to seven, or no anomalies found. This approach remedied 5,181 neurons and resolved the majority of over-packed stems, resulting in an inter-stem angle distribution in the small-angle range that closely aligns with that of EM-derived neurons (highlighted by red arrows in **Fig. 2e**). Consequently, the number of primary dendritic stems was considerably reduced, making it much similar to those in other datasets (**Fig. 2b**). When examining only neurons from the middle temporal gyrus (MTG), *i.e.*, the same brain region as for EM dataset, we observed more reduced distribution, with an upper boundary matching that of EM-derived neurons (**Fig. 2b**).

We verified the neuron structures of the remedied dataset by quantifying four features that assess tapering rates and branch orientations relative to the soma and parent branches (**Fig. 2f**). The results indicate that the ranges of all four features in ACT-H8K neurons fall within those of EM-derived neurons. Such statistics confirmed the consistency of both datasets.

### Reconstruction fidelity enables robust morphological quantification

To address the unique complexity of human tissue, we integrated deep-learning models, tailored enhancement and post-tracing EM spatial priors that render the updated protocol more powerful. Consequently, the pipeline achieved high fidelity for subsequent analyses (**Fig. S4a-b**). Compared to alternative approaches, LetsACT enhanced fiber integrity by reducing the number of breaks by 42.6% compared to the baseline, surpassing clDice (25.8%) and SkelRec (40.8%; **Fig. S4c**, left column). Additionally, LetsACT improved skeleton segmentation accuracy from 87.7% (baseline) to 94.3%, corresponding to a 6.6 percentage-point improvement (right column in **Fig. S4c**). We also assessed reconstructions using another three topological metrics, i.e., OPT-J, OPT-P, and OPT-G (Citraro et al., 2020), which evaluate consistency with manual annotations through node matching, path matching, and subgraph matching, respectively. LetsACT achieved F1 scores of 0.917, 0.954, and 0.864 across the three metrics (**Fig. S4d**). Reconstructions generated by LetsACT achieved the highest accuracy for three critical L-Measure features, i.e., “Number of Branches,” “Number of Tips,” and “Total Length”. Finally, our approach reduced the discrepancies to “gold standards” in relative feature values by 18.5%, 16.8%, and 22.1%, respectively, compared to the baseline, outperforming all competing methods (**Fig. S4e**).

We also compared our reconstructions to manually annotated “gold standards” on the test set (*n*=232). Quantitative evaluation was conducted across 11 morphological features defined by L-Measure (Scorcioni et al., 2008), a standardized repertoire critical for assessing reconstruction fidelity. The ratios of automated to manual measurements for each feature are presented as violin and box plots (**Fig. S5a**). Most features exhibited median ratios ranging from 0.9 to 1.0. In contrast, features such as total neurite length showed modest deviations below 1, suggesting minor overall under-tracing by the automated approach (**Fig. S5a**).

We next presented side-by-side comparisons of representative neurons for different ratios for “Number of branches” and “Total length” (**Fig. S5b**). These examples highlight that automated reconstructions effectively capture overall morphology, with discrepancies confined to distal branches at lower ratios. Additionally, comparison of these features between manual and automated annotations revealed a strong resemblance in their correlation matrices (**Fig. S5c-d**), indicating that the automated reconstruction platform achieves reasonable fidelity while supporting scalability for large-scale morphology datasets.

### Soma and dendritic branches of pyramidal cells shrink in infiltrated tissues

The relationship between neuronal morphology and brain disease represents a remarkable and rapidly developing field of study. For instance, in TDP-43-associated Parkinson’s disease dementia and temporal lobe degeneration, affected neurons show morphological degeneration (Armstrong et al., 2010, 2014). Similarly, neurons from patients with Kleefstra syndrome and autism show reduced length and fiber density (Nagy et al., 2017). Inspired by these observations, we investigated morphological differences between pyramidal and nonpyramidal cells across normal (*n*=8) and tumor-infiltrated (*n*=5) tissues from single-center glioma cohorts (**Fig. 3a**). Note that “normal” refers to tissue samples obtained along the surgical incision path that showed no evidence of tumor infiltration, whereas “tumor infiltrated” tissue was identified based on pathological assessment of tumor invasion.

**Fig. 3.**
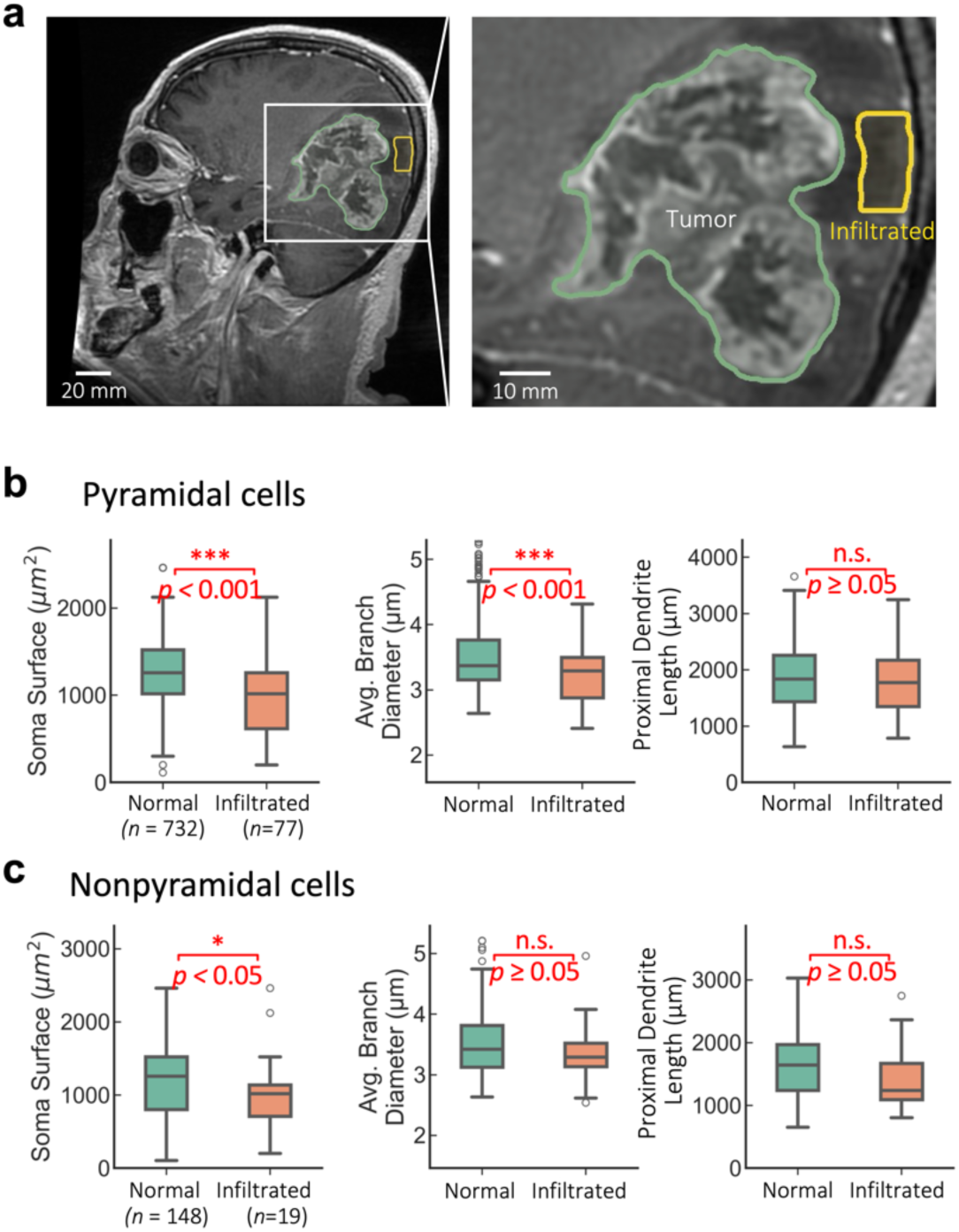
Morphological differences between neurons in normal versus tumor-infiltrated tissues. **a,** An example of a surgeon’s annotation of tumor-infiltration (yellow) on an MRI image. The remaining gray tissues are normal tissues. **b,** Morphological variations (e.g., soma surface, branch caliber, and proximal dendrite length) between pyramidal cells from normal tissues (*n*=732) and that from infiltrated tissues (*n*=77), with statistical significance labeled: n.s. (*p* > 0.05); \**p* < 0.05; \*\**p* < 0.01; \*\*\**p* < 0.001, based on two-sided Mann-Whitney U tests. **c,** Morphological variations for non-pyramidal cells. Statistical significance of differences between normal cells (*n*=148) and tumor-infiltrated cells (*n*=19).

Compared with neurons from normal tissue, those in tumor-infiltrated pyramidal cells exhibited reductions in soma surface area (*p* < 0.001) and branch caliber (“Average Branch Diameter”; *p* < 0.001; two-sided Mann–Whitney U test; **Fig. 3b**). In contrast, proximal dendrite length (dendritic length within a 100-µm radial distance from the soma) was indistinguishable (*p* ≥ 0.05; **Fig. 3b**). These patterns were confirmed by one-way ANOVA test (*p* = 7.76×10⁻⁴, 2.71×10⁻^4^, and 3.19×10⁻^1^, respectively). Nonpyramidal cells exhibited a gentler response to tumor infiltration. In specific, soma surface area and branch caliber were marginally reduced, while proximal dendrite length remained stable (**Fig. 3c**). However, it is important to note that the sample size for infiltrated nonpyramidal cells in this dataset was limited (*n*=19). Expanding the cohort in future studies may be necessary to achieve more robust statistical conclusions.

To ensure that these conclusions were not driven by technical biases, we performed multiple complementary robustness checks. (1) All tissue slices were collected from the same hospital and processed using identical fixation, staining, and imaging protocols, and reconstructions were performed by operators blinded to tissue condition; (2) When the analysis was restricted to neurons sampled from matched cortical regions (parieto-occipital junction), morphological changes persisted for soma surface and branch caliber (*p* < 0.001), and proximal dendrite length showed no difference (**Fig. S6a**); (3) We further evaluated statistical robustness by averaging the results of 10 trials, in which a random 50% subset of normal neurons was compared against the infiltrated neurons. The previous morphological changes remained (**Fig. S6b**); (4) Given that pyramidal and nonpyramidal cells were sampled from overlapping tissue sets (8 shared tissues; total *n*=12 and *n*=9, respectively), the morphological changes specifically observed in pyramidal cells is unlikely to result from systematic biases. These consistent findings across patients, cortical regions, and neuron subsets indicate that the observed somatic and branch changes is a robust phenomenon specifically associated with tumor infiltration rather than confounding technical or biological variables.

These results indicate a cell-type-specific morphological vulnerability in the human glioma-infiltrated cortex. Pyramidal neurons undergo consistent shrinkage in both soma surface area and branches, but nonpyramidal neurons may not. This difference may arise from distinct metabolic demands, synaptic partnerships, or molecular responses to tumor-derived factors. Functionally, the pronounced degeneration of pyramidal neurons could disrupt the excitatory backbone of local circuits, contribute to seizure susceptibility, and exacerbate cognitive deficits. Our findings underscore neuronal morphology as a sensitive indicator of the peritumoral microenvironment and support further investigation of glioma–neuron interactions and neuroprotective strategies to preserve pyramidal neuron integrity.

### Somatic volume reduction and increased anisotropy of pyramidal cells in infiltrated tissues

As the somatic surface area of pyramidal cells decrease in infiltrated tissues (**Fig. 3**), we also investigated the morphological deformations of nearby neurite compartments. 3-D segmentation of neuronal somas revealed clear morphological changes in excitatory pyramidal cells, whereas nonpyramidal somas remained largely unaffected (**Fig. 4**).

**Fig. 4.**
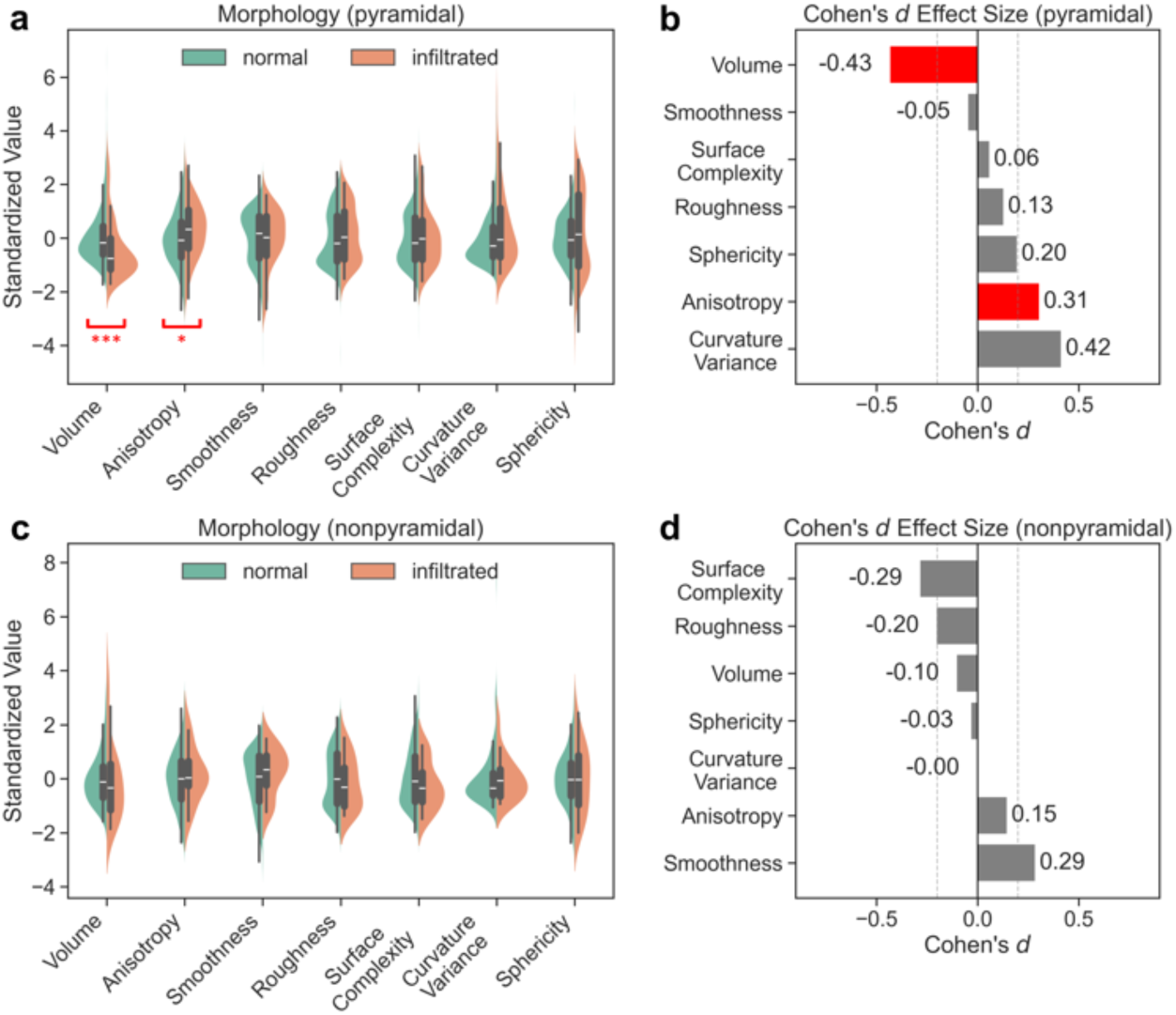
Somas of pyramidal cells exhibit morphological deformations in infiltrated tissues. **a**, Violin plots showing seven standardized 3D soma shape descriptors for pyramidal neurons from control (normal, green; *n* = 728) and tumor-infiltrated (orange; *n* = 77) human cortical tissue. Overlaid box plots indicate the 25th and 75th percentiles (box edges), median (central line), and whiskers extending to 1.5 × the interquartile range. **b**, Cohen’s *d* effect sizes for pyramidal neurons. Red bars highlight features with large, statistically significant changes (two-sided Mann-Whitney U test \**p* < 0.05; \*\*\**p* < 0.001); gray bars indicate non-significant or small effects. The dashed vertical line marks |*d*| = 0.2 (conventional threshold for a small effect). **c**, Same as (**a**) but for nonpyramidal neurons (normal: *n* = 147; infiltration: *n* = 18). **d**, Cohen’s *d* effect sizes for nonpyramidal neurons.

In pyramidal cells, tumor exposure induced noteworthy shifts across independent shape descriptors (**Fig. 4a,b**). Soma volume was notably decreased (Cohen’s *d* = -0.43, *p* < 0.001), consistent with the reduced soma surface area previously observed (**Fig. 3c**). The curvature variation showed the second-largest effect (*d* = 0.42), together indicating that tumor-exposed pyramidal somas are generally smaller and adopt more convoluted contours. This potentially reflects a loss of the subtle local invaginations and regularities that characterize healthy pyramidal cells. Anisotropy was also modestly increased (*d* = 0.31, *p* < 0.05). In contrast, nonpyramidal cells exhibited no comparable variation (**Fig. 4c,d**). None of the seven descriptors reached statistical significance or a meaningful effect size (|*d*| > 0.4).

Our results show that somas statistically exhibit not only volumetric shrinkage but also additional shape alterations, including increased anisotropy and greater curvature variance, which likely reflect underlying cytoskeletal reorganization. These somatic deformations occur noticeably in excitatory pyramidal neurons. This might suggest an unexpected cell-type specificity that the presence of the tumor may impose differential mechanical or biochemical pressures on excitatory and inhibitory neuronal populations.

### Pyramidal cell shrinkage attenuates from proximal to distal in infiltrated tissues

As somas and dendritic arborization shrink in infiltrated pyramidal cells (**Fig. 3-4**), we next investigated whether the observed changes were uniform or represented an independent degenerative phenotype. To test this, we normalized morphological features, including coordinates and radii, of each neuron to its soma size (**Fig. 5b**, **Fig. 5c**).

**Figure 5.**
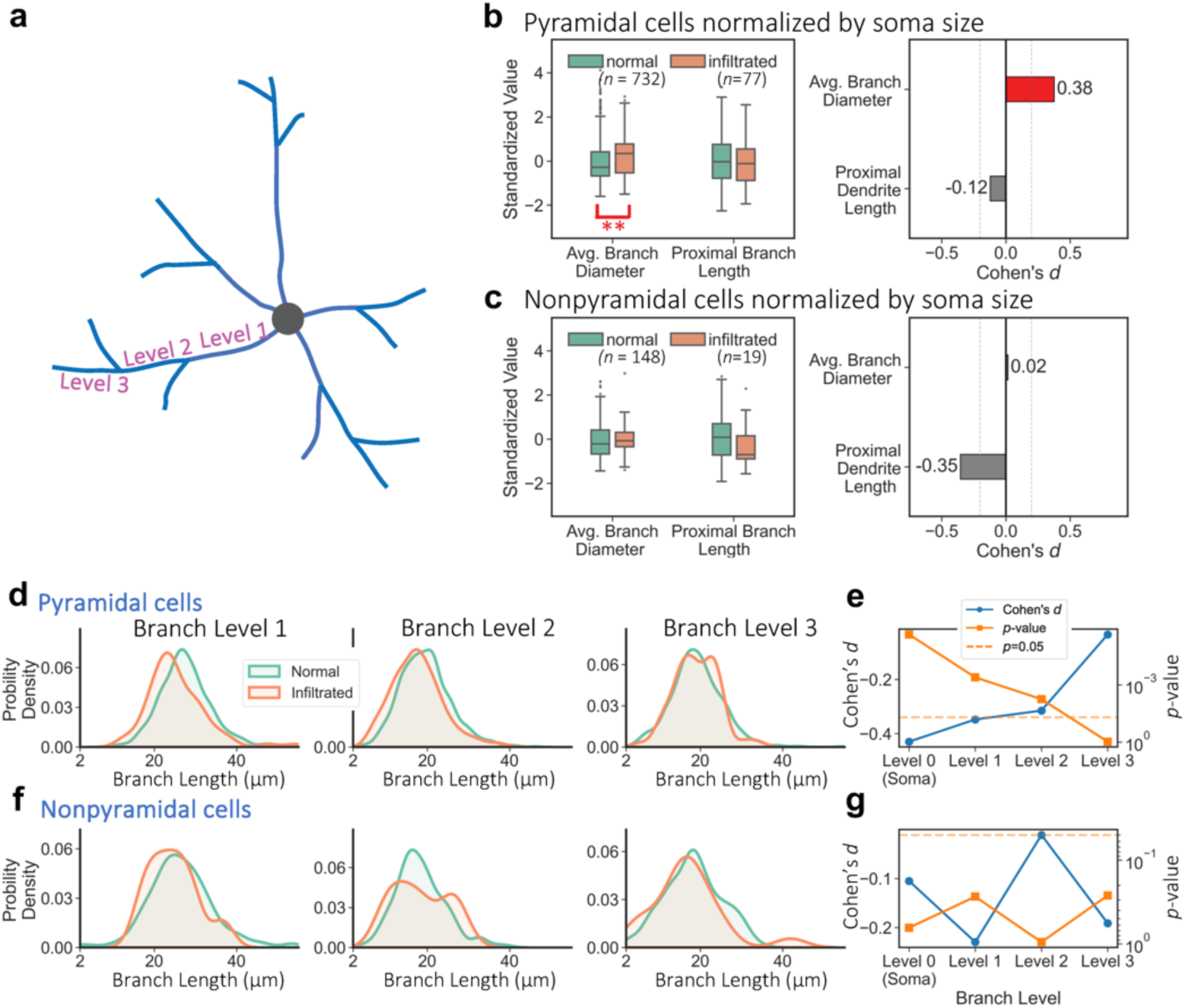
Attenuating shrinks of pyramidal cells in infiltrated tissues. **a**, Schematic illustration of branch levels. **b,** Morphological comparison of pyramidal (**b**) and nonpyramidal (**c**) neurons after normalization by soma size. **Left:** Split box plots display the standardized distributions of Average Branch Diameter and proximal dendrite length for normal (green) and infiltrated (orange) neurons. Statistical significance is indicated (\*\**p* < 0.01). **Right:** Effect size analysis using Cohen’s *d*. Pyramidal neurons show a marked difference in average branch diameter (red bar; *d*=0.38), whereas proximal dendrite length and nonpyramidal features show negligible effect sizes (grey bars). **d,** Association of branch length reduction and branch levels for pyramidal cells. Probability density plots comparing the distributions of non-terminal branch length in original un-normalized morphologies across branch levels 1, 2, and 3. **e**, Quantification of morphological changes across hierarchical levels. Effect sizes from level 0 (soma) to level 3 (distal) are shown in blue, and *p*-values are plotted in orange. The dashed orange line indicates the statistical significance threshold (*p*=0.05). **f, g**, Analysis analogous to **d** and **e** for nonpyramidal cells.

Pyramidal cells showed a non-uniform morphological change between somas and dendritic arborizations. While the proximal dendritic neurite length in normalized morphologies had negligible differences between normal and infiltrated tissues (Cohen’s *d* = -0.12), the reduction in average branch diameter was reversed (Cohen’s *d* = 0.38; **Fig. 5b**). This reversal indicates that the shrinkage of dendritic branches in infiltrated pyramidal neurons is not simply a passive geometric scaling commensurate with cell body shrinkage, but rather suggests a relative resilience of the neurite shaft. In contrast, nonpyramidal neurons displayed high stability, and normalizing by soma size yielded negligible effect sizes for both average diameter (Cohen’s *d* = 0.14) and proximal dendrite length (Cohen’s *d* = -0.35; **Fig. 5c**).

To identify the subtle pattern of the morphology shrinkage, we analyzed these changes of branching hierarchies using original, non-normalized data (branch levels 1–3; **Fig. 5d**). Generally, branch lengths in infiltrated pyramidal cells exhibited a modest reduction (**Fig. 5d**), with Cohen’s *d* values ranging from - 0.35 to -0.03 (level 1 to level 3; **Fig. 5e**). Accordingly, Mann-Whitney *p*-values increased from level 1 (4.08×10^-4^) to level 3 (0.96), indicating that morphological changes attenuate from proximal to distal compartments. Comparative analysis revealed that the soma underwent the most prominent change (Cohen’s *d*=-0.43, *p*=2.3×10^-6^; **Fig. 5e**). In contrast, nonpyramidal cells showed negligible effect sizes and larger *p*-values (**Fig. 5f**) for both somas and dendritic branches, and the monotonic spatial transition observed in pyramidal populations was not found (**Fig. 5g**). Given that nonpyramidal cells share the same patient sources, the heterogeneous changes observed in pyramidal cells underscore a non-uniform susceptibility to the peritumoral microenvironment.

In summary, infiltrated pyramidal neurons exhibit a distance dependent morphological phenotype. The structural deformation is characterized by marked somatic shrinkage that gradually lessens along the dendritic arbor. This nonuniform pattern indicates that the somas and nearby dendritic branches are disproportionately vulnerable to oncogenic stress.

### Infiltrated cells may undergo a defensive morpho-genetic response during oncogenesis

Given that infiltrated pyramidal cells exhibit heterogeneous changes in soma morphology (**Fig. 3**, **Fig. 4**) and branch diameter (**Fig. 3**), a critical question arises (**Fig. 6a**): do these morphological changes represent a snapshot toward apoptosis (cell death), or an adaptive “hunkering down” to defend against oncogenic stress? This question cannot be resolved using morphological data alone. To address this, we integrated our morphological reconstructions with both the in-house 10× Visium spatial transcriptomic data and a public CGGA dataset of bulk mRNA sequencing (Bao et al., 2014; Wang et al., 2015; Zhang et al., 2022; Zhao et al., 2017, 2021). This integrative approach considers several stages of oncogenesis, allowing a joint analysis of phenotypes and genotypes to infer a potential mechanism that drives neuronal changes (**Fig. 6b**).

**Fig. 6.**
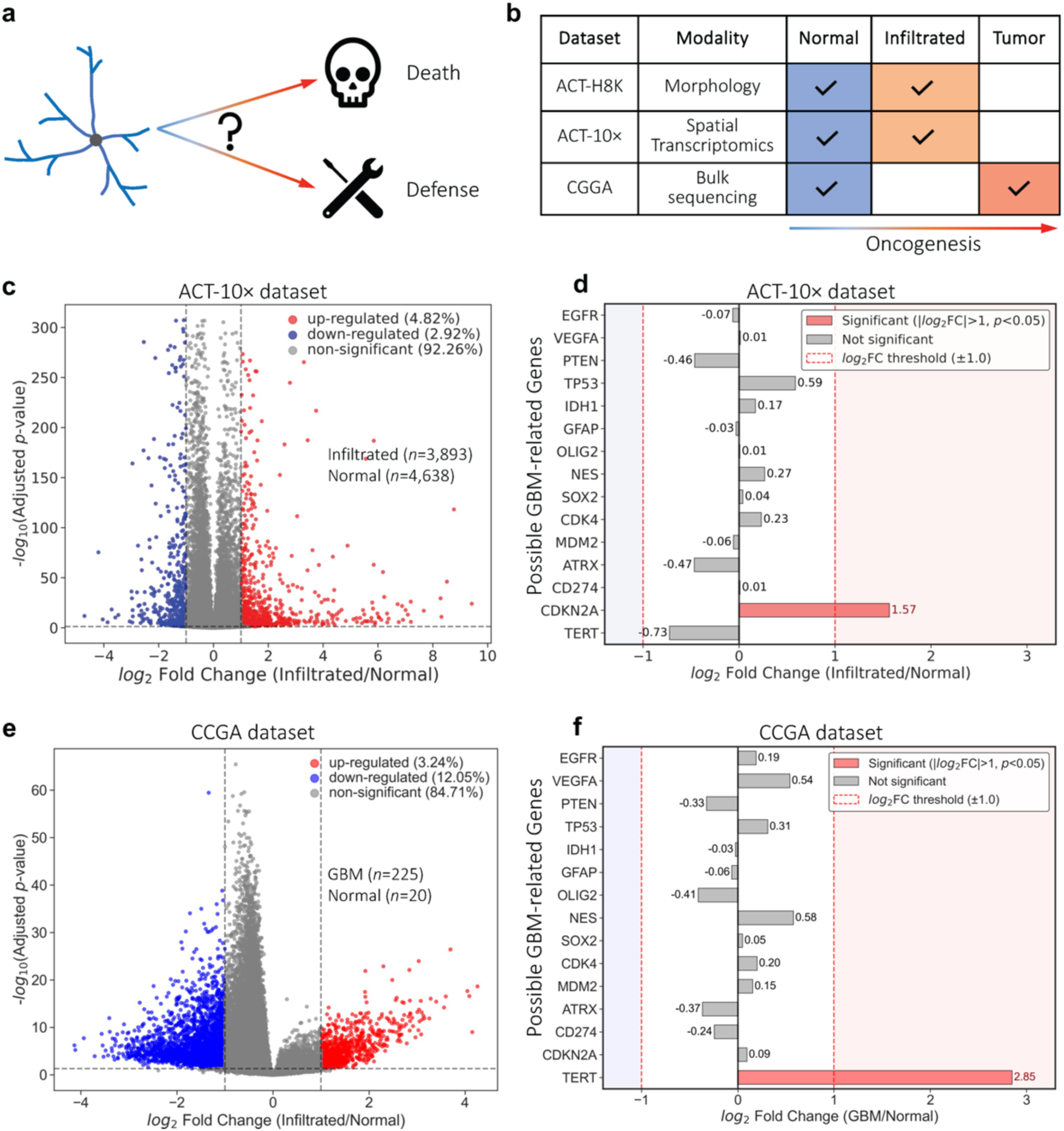
Integrative phenotype-genotype modeling reveals a potential mechanism of neuronal defense against glioblastoma. **a**, Schematic illustration of two possible morphological reaction models during oncogenesis. **b**, The three datasets involved. The composite dataset compromise datasets of different tumor stages and modalities. Tick marks and shading indicate the presence of the corresponding stage. **c**, Volcano plot illustrating the global differential gene expression between infiltrated spots (*n*=3,893, P66) and normal cell controls (*n*=4,638, P65) from the junction area of parieto-occipital cortex. Each dot represents a single gene mapped by its magnitude of change (log_2_FC, *x*-axis) and statistical significance (-log_10_ adjusted *p*-value, *y*-axis). Red points indicate significantly up-regulated genes (log_2_FC > 1, *p*_adj_ < 0.05), and blue points indicate significantly down-regulated genes (log_2_FC < -1, *p*_adj_ < 0.05). The vertical dashed lines denote the 2-fold change threshold, and the horizontal dashed line marks the significance cutoff (*p*_adj_ < 0.05). **d**, Differential expression analysis of selected canonical glioma-related genes. Bars represent the log_2_FC for each gene. Only *DKN2A* (red bar) exhibited up-regulation exceeding the predefined fold-change threshold (|log_2_FC| > 1, *p*_adj_ < 0.05). Gray bars indicate genes that did not meet the fold-change criteria and statistics in this cohort. **e-f**, Similar volcano plot (**e**) and bar plot (**f**) for GBM samples (*n*=225) and normal tissue controls (*n*=20) from the CGGA repository.

Fistly, we performed a differential expression analysis comparing infiltrated 10× Visium spots (*n*=3,893) against normal spots (*n*=4,638) from the same junction area of parieto-occipital cortex. The global expression profile revealed widespread transcriptomic dysregulation. We identified a distinct set of up-regulated (red dots, 4.82%) and down-regulated genes (blue dots, 2.92%, **Fig. 6c**) using a stringent threshold of |log_2_ Fold Change (FC)|>1.0 and adjusted *p*<0.05. We also interrogated a panel of canonical glioblastoma biomarkers to validate their expression status (**Fig. 6d**). The critical tumor suppressor gene *CDKN2A* was overexpressed in infiltrated tissues, exhibiting a high log_2_FC of 1.57 (**Fig. 6d**). Another common tumor suppressor, *TP53*, also showed overexpression (log_2_FC=0.59, **Fig. 6d**). In contrast, tumor-promoting genes such as *TERT* and *ATRX* were downregulated (**Fig. 6d**). In both infiltrated and normal tissues, *TERT* expression is three orders of magnitude lower than *CDKN2A* expression. The upregulation of tumor suppressors and downregulation of tumor promoters suggests that infiltrated cells may be undergo a defensive response against the tumor microenvironment, possibly mediated through *CDKN2A*.

We also benchmarked bulk mRNA sequencing of GBM tissues (*n*=225) against non-glioma controls (*n*=20, **Fig. 6e**). Gene dysregulation was more extensive in GBM tissues (15.29%, **Fig. 6e**) compared to infiltrated cortical spots (7.74%, **Fig. 6c**). Specifically, GBM tissues exhibited a higher proportion of downregulated genes (12.05% vs. 2.92% in infiltrated tissues) but a lower proportion of upregulated genes (3.24% vs. 4.82%; **Fig. 6c, e**). Regarding canonical GBM-related gene signatures, the most prominent change in the tumor was the overexpression of *TERT* (log_2_FC =2.85, *p*<0.05; **Fig. 6f**), consistent with its role in telomere maintenance and immortalization in aggressive gliomas. Additionally, the upregulation of classical GBM markers such as *VEGFA* and *NES* was more pronounced in the bulk tumor data (**Fig. 6f**) compared to the infiltrated tissue (**Fig. 6d**).

An interesting feature of the differential expression landscape lies in the joint distribution of fold change (log_2_FC) and statistical significance (-log_10_*p*). In bulk GBM tissues, the upregulated fraction (positive log_2_FC) exhibited a tightly packed, linear correlation (**Fig. 6e**, right half of the panel), reflecting a consistent signal from shared tumor drivers across samples. Conversely, the downregulated fraction displayed a dispersed distribution (**Fig. 6e**, left half of the panel). In the transcriptomic analysis of infiltrated versus normal spots, we observed high dispersion across both upregulated and downregulated domains (**Fig. 6c**). This “scattered” topology may be attributed to the heterogeneity of the cellular response to infiltration, or the convolution of various cell types.

To determine if the dispersed pattern in the infiltrated cells (**Fig. 6c**) was driven by cellular heterogeneity, we performed a stratified analysis restricted to spots dominated by a single cell type (ranking 1st in abundance and constituting >40% abundance probability of a spot). We found that excitatory upper-layer intratelencephalic neurons in infiltrated tissues exhibited a relatively stable profile with low level of dysregulation (4%, **Fig. S7a**). In contrast, glial populations (astrocytes and oligodendrocytes) showed high reactive states, characterized by larger proportion of dysregulated genes (7.81% for astrocytes and 11.64% for oligodendrocytes, **Fig. S7c,e**). Due to the extremely low expression of *TERT* within individual cell types, reliable statistical analysis of this gene was not feasible (**Fig. S7b,d,f**).

These data propose a potential morphological and genetic response model for GBM. While the core GBM genotype is defined by massive gene repression and consistent driver activation (e.g., *TERT*), the infiltrated cortex is characterized by a heterogeneous, predominantly upregulated profile consistent with host tissue stress or defense mechanisms (e.g., *CDKN2A*). This molecular signature parallels the morphological changes observed in dendritic arborization and soma geometry. Together, these findings imply that the peritumoral microenvironment may involve a selective, adaptive remodeling of phenotype and genotype in response to oncogenic stress.

## Discussion

The complexity of human neuronal image necessitates tailored algorithmic designs. As a critical step toward digitizing the human brain at single-cell resolution, we developed an automated, AI-driven platform capable of generating large-scale, high-fidelity reconstructions. By integrating this platform with a high-throughput tomography pipeline, we successfully reconstructed 8,398 neurons across diverse regions of the human brain. These reconstructions illustrate the dataset’s potential for quantifying morphological changes associated with pathology, such as tumor infiltration. We release this resource to the neuroscience community with the hope to help accelerate the exploration of human brain anatomy and functions.

A key innovation of the LetsACT approach is its capacity for multimodal data integration. This is especially important because different laboratories frequently use distinct experimental practices, which results in large datasets that are difficult to combine in a unified analysis. By incorporating cutting edge datasets from both light microscopy and electron microscopy, we were able to substantially improve our initial light microscopy reconstructions. This integration reduced reconstruction artifacts such as artificial over branching and improved overall structural fidelity.

We further advanced morphogenetic integration by linking morphological alterations to dysregulation of glioblastoma related genes in both our in house spatial transcriptomic data and publicly available bulk sequencing datasets. We observed an inverse expression pattern: infiltrated tissues showed upregulation of tumor suppressor genes and downregulation of tumor promoter genes, whereas tumor tissues showed the opposite trend. These observations support the possibility that the morphogenetic changes we report may reflect an adaptive defensive response to tumor invasion rather than a simple trajectory toward apoptotic degeneration.

Our analyses reveal morphological alterations, such as differing vulnerability between pyramidal and nonpyramidal neurons in glioma infiltrated tissue. However, the molecular and cellular mechanisms responsible for these changes would be interesting for a future study. The central aim of this work is to establish a scalable framework for high throughput generation of human neuron data with demonstrated applications for their use for meaningful findings. Our biological discoveries illustrate the potential of our approach and dataset to guide future investigations into metabolic, synaptic, or genetic processes associated with the observed structural changes.

As with all investigations utilizing human surgical resections, our study could be further improved when larger datasets become availability. The number of reconstructed tumor-infiltrated neurons remains modest (*n*=77 pyramidal cells) compared to normal controls (*n*=732). While these constraints are inherent to clinical neuroscience, the present dataset that spanning 13 patients and over 800 pyramidal neurons from the same hospital and staining protocol represents one of the largest cohorts of its kind. The consistency of morphological changes observed across patients, cortical regions, and multiple validation tests supports the robustness of our findings. We are actively expanding this dataset and incorporating additional pathological controls to validate these results in future work.

The infiltrated region of glioma presents a multifaceted pathological environment. This complexity arises from the heterogeneity of the cellular composition, which spans diverse brain regions and includes distinct cell types, such as neurons of various neurotransmitter phenotypes, glia, and infiltrating blood cells. Concurrently, the specific tumor microenvironment, such as high cellular density and intense survival pressure, further complicates the exploration of underlying mechanisms. Given this intricate biological context, single-modality approaches often provide an incomplete view. By integrating light microscopy, electron microscopy, and transcriptomic data across various tumor states, our study offers an exploratory framework designed to navigate this complexity. This multimodal approach serves to deconvolute the tissue microenvironment, facilitating the distinction of host adaptive responses from the broader context of tumor invasion.

Despite methodological advances, mapping whole-brain connectivity, particularly the reconstruction of distal axons, remains a formidable challenge. While we observed that secondary immunohistochemical staining enhances neuronal completeness (**Fig. S8**), capturing entire axonal projections is often precluded by the physical limitations of tissue slicing and the ethical constraints limiting *in vivo* viral tracing in humans. Nevertheless, our previous research in the mouse hippocampus demonstrated that local dendritic morphologies, when analyzed via microenvironment representations, are predictive of long-range projection patterns (Liu et al., 2025). This suggests that even in the absence of full axonal reconstruction, detailed dendritic analysis may offer a valuable proxy for inferring functional connectivity.

## Methods

### Ontology and Nomenclature for Brain Regions

During surgical procedures, we captured at least two photographs per sample: an intraoperative image documenting the sampling location within the brain, and a postoperative image annotated by physicians annotating location, brain region, and anatomical orientation. Physicians manually segmented MRI images from corresponding patients, marking 3D locations of tumors and sampled tissues. To standardize the heterogeneous brain region labels used across institutions and physicians, we mapped masks to the Yale Brain Atlas (McGrath et al., 2022). Detailed surgical and imaging protocols are described in (Han et al., 2023). The studies involving human tissue were approved by the ethics committees of Beijing Tiantan Hospital, The First Affiliated Hospital of Nanjing Medical University, Eastern Theater Command General Hospital, and Huashan Hospital, with informed consent obtained from all participants.

### ACT-Specific Enhancement

#### Halo noise modeling

We modeled perisomatic halo artifacts using a Gaussian diffusion process, based on the observation that halo intensity attenuates gradually with distance from the soma.

First, we extracted high-intensity regions *I*_*hi*_ by thresholding voxels above the 90th percentile of the original image *I*. We then applied Gaussian filtering to generate a simulated diffusive halo:

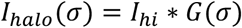

where *G*(*σ*) denotes a Gaussian kernel with a standard deviation of *σ* and ∗ represents convolution.

We combined *I*_*hi*_ with the simulated halo noise *I*_*halo*_(*σ*) to produce the composite image:

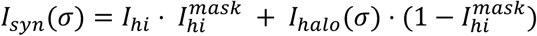

where 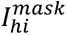 represents the mask estimated by *I*_*hi*_ > 0.

To determine the optimal Gaussian kernel, we maximized the Structural Similarity Index Measure (SSIM) between the original and composite images:

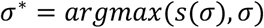

The resulting simulated noise regions was used to guide adaptive enhancement of distal neurites while preventing overexposure near somas.

#### Image enhancement

We normalized *I*_*syn*_ to generate a spatially varying gamma coefficient map:

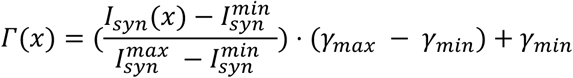

where 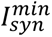 and 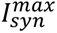 represent minimum and maximum voxel values, and *γ*_*min*_ = 0.5 and *γ*_*max*_ = 1.0 define the gamma correction range.

We performed spatially adaptive gamma transformation using *Γ*(*x*) as the local power coefficient:

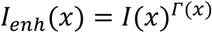

### Segmentation

#### Label generation

We generated segmentation labels by combining annotated morphologies with raw image data. First, we extracted potential neurite voxels by thresholding at the 50th percentile of voxel intensities. We retained only voxels within the connected component containing the target soma, yielding 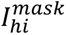. We then estimated radii for all skeletal nodes (excluding the soma region) and converted the skeleton to a volumetric mask (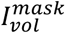). Simultaneously, we binarized the original image to create a smooth-edged foreground mask (*I*_*bin*_). To prevent discontinuities, we generated a skeletal mask (*I*_*skel*_) with a constant one-voxel radius. The final segmentation label was computed as:

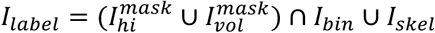

where ∩ and ∪ represent element-wise AND and OR operations, respectively.

We divided the 1,332 annotated neurons into a training-validation set (*n* = 1,100) for five-fold cross-validation and a test set (*n* = 232) for quantitative evaluation.

### Connectivity enhancing loss for tubular structures

We developed a connectivity-enhancing loss (CEL) to preserve neurite continuity:

1. False-Negative Skeleton (FNS): We computed the skeletonized ground truth (*I*_*skel*_) and identified false-negative regions: *I*_*fns*_ = *I*_*skel*_ ⋅ *ReLU*(0.5 − *I*_*output*_)
2. Loss calculation: CEL was computed as the mean intensity of false-negative skeleton voxels: 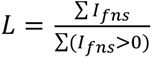

### Model training

We employed nnUNet (Isensee et al., 2021) for segmentation. Training proceeded in two phases: initial training from scratch using combined cross-entropy and Dice losses, followed by fine-tuning with additional CEL loss. We used stochastic gradient descent with an initial learning rate of 1×10^-2^, decaying exponentially to 3×10^-5^ during phase one. For fine-tuning, we set the initial learning rate to 1×10^-5^, with 0.9-fold decay every 100 epochs after epoch 150. All training utilized NVIDIA RTX 3090 GPUs.

### Evaluation Metrics

We proposed two customized metrics to assess segmentation integrity:

#### Number of Skeleton Breaks

We identified false-negative regions by comparing segmentation results with dilated skeletal masks, then counted connected components within these regions:

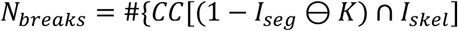

where ⊖ denotes dilation with a cross-shaped kernel (*K* = 3), and #{CC[⋅]} represents the number of connected components. Dilation prevents misclassification of minor centerline shifts as true breaks.

#### Skeleton Accuracy

We calculated the ratio of correctly segmented skeleton voxels:

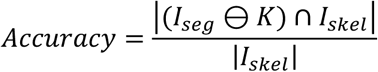

where |*I*| denotes the total number of positive voxels.

### Data Generation and Quality Control

We acquired 14,763 neuronal image stacks using ACTomography. Approximately 6,000 stacks were excluded based on the following criteria:

1. **Noisy soma imaging**: Stacks with improper soma visualization, including hollow somas or those with high diffusive, mushroom-shaped artifacts.
2. **Intermingled neurons**: Stacks containing multiple neurons in close proximity that could introduce topological errors.
3. **Poor imaging**: Stacks exhibiting tissue tears or high-intensity noise volumes that could cause over-tracing artifacts.

The remaining images underwent reconstruction using APP2 with default settings. Reconstructions falling within the lowest 5th percentile of total skeletal length were excluded. Retained reconstructions were subsequently refined using priors derived from the EM dataset (detailed below).

### EM-LM synergized refinement

#### Comparing datasets

We utilized three public datasets to compare primary stem distributions: (1) **DeKock dataset**: 89 pyramidal neurons from human middle temporal gyrus (MTG), containing dendritic structures from surgical tissues (Mohan et al., 2023). (2) **Allen dataset**: 297 human cortical neurons of diverse cell types from the Allen Human Cell Types database (Berg et al., 2021). (3) **H01 dataset**: Neuronal skeletons from C3 segment layers, and corrected using an LLM-based protocol for denoising and reconnection (Gu et al., unpublished data). Following quality control, 3,446 neurons were retained for comparative analysis and spatial distribution modeling.

#### Gaussian Mixture Model for neuron stems

We modeled the spatial distribution of primary stems using GMM. For each of 14,790 primary stems in the H01 dataset, we calculated the number of neighboring stems within angular ranges of 10°, 20°, 30°, 40°, and 50°, generating a 5-dimensional angular-distribution representation (ADR) that captured local spatial arrangements. We selected an 11-component GMM based on Bayesian Information Criterion (BIC).

#### Refinement

To correct topological artifacts in the initial ACT-H8K dataset, we implemented an iterative EM-LM synergized refinement process. The GMM identified anomalous stems, which were then merged with spatially closest neighbors using an iterative detection-and-merging algorithm. This process continued until either the stem count per neuron was reduced to seven or no additional anomalies were detected.

### Morphological Features

We computed 22 morphological features using Vaa3D (Peng et al., 2014), with detailed descriptions available in (Liu et al., 2024). For the comparative analyses between infiltrated and normal neurons, we modified the “Average Diameter” calculation to exclude compartments within the soma range (denoted as “Average Branch Diameter”), thereby decoupling this metric from “Soma Surface”.

### Spatial-Transcriptomics

#### 10× Visium data processing

We acquired spatial transcriptomic data using 10× Visium CytAssist with a 6.5 × 6.5 mm capture area. Fresh-frozen OCT-embedded brain samples with RNA Integrity Number (RIN) > 4.0 were processed through hematoxylin and eosin (H&E) staining, probe hybridization, library construction, and Illumina sequencing. Raw FASTQ files and histology images were processed per sample using Space Ranger software (v2.0.1) with the GRCh38-2020-A reference genome and Human Transcriptome v2 probe set.

Cell-type composition was estimated for each spot using *cell2location* (Kleshchevnikov et al., 2022), by combining single-cell transcriptomic data from adult human brain (Siletti et al., 2023) with our spatial transcriptomics data. Only cells from corresponding brain regions (and Brodmann areas 5-7 and 19 for P65 and P66), were used for estimation. The initial gene set was filtered using scanpy.filter_genes (*cell_count_cutoff* = 5, *cell_percentage_cutoff2* = 0.15, *nonz_mean_cutoff* = 2). The cell2location model was trained for 200 epochs on the single-cell dataset, with key prediction parameters *N_cells_per_location* = 8 and *detection_alpha* = 20.

### Soma Mask Estimation

To isolate the soma from the neurite segmentation maps, we developed an automated 3D morphological approach. First, to maximize computational efficiency, the segmentation volume was cropped to the bounding box of the non-zero mask with a padded margin. Because the original imaging data was anisotropic, the cropped volume was resampled to an isotropic resolution based on the physical pixel dimensions (*z*/*xy* ratio) to ensure consistent morphological operations. Binary hole-filling was applied to the isotropic mask to ensure the soma volume was solid.

The soma was defined as the largest compact structure within the segmentation. We employed an iterative morphological opening process using spherical structuring elements with incrementally increasing radii (ranging from 2 to 15 voxels). This operation progressively eroded thinner neurite branches while preserving the thicker soma structure. At each iteration, only the largest connected component of the remaining mask was retained to eliminate disconnected fragments.

To determine the optimal morphological kernel size, we fitted a bounding ellipsoid to the remaining foreground voxels at each iteration using Principal Component Analysis (PCA) on the voxel coordinates. The center, orientation (eigenvectors), and semi-axes were derived from the covariance matrix of the point cloud. We defined a “foreground ratio” metric, calculated as the proportion of the fitted ellipsoid’s volume occupied by the segmentation mask. The iteration terminated when this ratio exceeded a threshold of 0.65, indicating that the remaining structure corresponded to a compact, soma-like volume. The final soma mask was generated using the optimized kernel radius and transformed back to the original coordinate space.

To validate, we randomly selected a subset of 35 neurons and superimposed the generated masks onto the original image data on three orthogonal planes (*xy*, *xz*, and *yz*). Visual inspection confirmed a high degree of concordance between the segmentation boundaries and the somatic signal intensity (**Fig. S9**).

### Soma Morphological Feature Extraction

We computed a comprehensive set of morphological features to characterize the geometric properties of each cell. The soma volume was calculated as the product of the total foreground voxel count and the cubic physical resolution (*V* = *N_voxels_* × *resolution_xy_*^3^). To describe the global shape anisotropy, we utilized the semi-axes lengths (*r_x_*, *r_y_*, *r_z_*) derived from the ellipsoid fitting step. Based on these dimensions, we calculated the anisotropy as 1 - *r_min_*/*r_max_*), where are minimal and maximal values in (*r_x_*, *r_y_*, *r_z_*).

Sphericity was computed using the standard Wadell equation 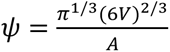, relating the soma volume (*V*) to its surface area (*A*). To quantify surface smoothness, we estimated local surface normal vectors using central differences within a 3×3×3 neighborhood. A smoothness score was derived from the consistency of normal orientations (dot products) between adjacent surface points, while surface roughness was defined by the variance of these orientations. Surface complexity was defined as the complement of the smoothness score (1 - *smoothness*). Additionally, local surface curvature was approximated by sampling surface patches (7×7×7 voxels) and performing a local spherical fit, where the curvature variance was estimated from the variance of voxel distances to the patch centroid.

### Morphological Normalization and Cropping

To test if neurite arborization patterns independent of soma size differences between cell types (pyramidal vs. nonpyramidal) and tissue environments (normal vs. infiltration), we applied a linear size normalization. A scaling factor was calculated for each experimental group relative to a reference baseline (pyramidal cells in normal tissue). This factor was defined as the ratio of the group-specific soma size to the reference size (*scale* = *S_group_* / *S_ref_*), where *S_group_* and *S_ref_* are the average size for the target group and reference group respectively. The 3D coordinates (*x*, *y*, *z*) and radii (*r*) of all neuronal reconstructions were normalized by dividing by this scaling factor, effectively mapping all neurons to the dimensions of the reference population.

Following normalization, the reconstructions were spatially cropped to enable fair comparation across neurons. In specific, we retained only those neurite segments situated within a spherical region of interest (ROI) with a radius of 100 μm centered on the soma. Segments extending beyond this boundary were discarded to ensure the analysis focused on the local connectivity potential of the neuron.

### Differential Expression Analysis

To identify transcriptomic differentiation associated with Glioblastoma (GBM), we performed a differential expression analysis comparing GBM samples against normal brain tissue controls. Gene expression values (FPKM) were download from the Chinese Glioma Genome Atlas (CGGA; https://www.cgga.org.cn/download.jsp) and then pre-processed to filter out low-abundance transcripts; genes were retained only if they exhibited an FPKM value > 0.1 in at least 10% of the samples. To account for systematic differences in library size and composition, a median-based normalization was applied to align the distribution centers of the GBM and normal cohorts.

For the assessment of differential expression, we calculated the log_2_ Fold Change (log_2_FC) for each gene as the base-2 logarithm of the ratio between the arithmetic mean expression of the GBM group and that of the normal control group. Statistical significance was evaluated using Welch’s t-test to account for unequal variances between the tumor and normal populations. To control for the false discovery rate (FDR) associated with multiple hypothesis testing, raw p-values were adjusted using the Benjamini-Hochberg (BH) procedure.

Differentially expressed genes (DEGs) were identified based on dual thresholds: a magnitude of change of |log_2_FC| > 1.0 (representing a >2-fold difference) and a statistical significance threshold of adjusted *p* < 0.05. Genes satisfying these criteria were classified as up-regulated or down-regulated, respectively. All analyses were implemented using custom scripts in Python.

## Data availability

The auto-reconstructed morphologies and 1,342 manually annotated reconstructions are available on Zenodo (doi: 10.5281/zenodo.15189542).

## Code availability

Source code for neuron segmentation and reconstruction is publicly available on GitHub at https://github.com/SEU-ALLEN-codebase/neuron_seg_human. Analytical scripts can be accessed at https://github.com/SEU-ALLEN-codebase/HumanMorphoMap. These projects utilize our in-house analytical library for neuron morphology, *pylib*, available at https://github.com/SEU-ALLEN-codebase/pylib. Other key dependencies include: Vaa3D-x (v1.1.4), scanpy (v1.11.2), cell2location (v0.1.4), PyTorch (v2.1.2+cu118 for neuron segmentation; v2.7.1+cu12.6 for cell2location).

## Acknowledgement

This project was mainly supported by several grants awarded to H.P. H.P. is a New Cornerstone Investigator and a SANS Senior Investigator. Y.L. is also supported by a start-up package offered by Fudan University. Additional support was provided by the Natural Science Foundation of Jiangsu Province (Grant No. BK20243064), the Fundamental Research Funds for the Central Universities (No. 2242023K5005), Project of Jiangsu Science and Technology Plan (No. BK20243058), National Science and Technology Major Project (No. 2025ZD0215100), NSFC 82272116, Science and Technology Commission of Shanghai municipality frontier innovation program (No. 24DP3200600), and Jiangsu Province Capability Improvement Project through Science, Technology and Education (No. ZDXK202225). We thank Xiaoqin Gu, Haoyu Zeng, Xiaoxuan Jiang, Lijun Wang, Shengdian Jiang, Lijuan Liu, Xin Chen, Ting Peng, Xinzheng Sun, and several other members in Peng Lab for assistance of data pipeline. We also thank Ed Lein, Javier DeFelipe, Ruth Benavides-Piccione, and Giorgio Ascoli for discussion.

## Author contributions

H.P. conceived, designed and supervised the project. H.P. contributed key ideas of this study, provided both experimental and analytical platforms, and instructed the detailed analyses. Y.L. executed the study along with the help of all co-first authors, and contributed especially on the algorithms. Z.Y. conducted brain registration and also contributed to data analyses. L.Z. contributed to the key components of the cell injection system. W.Y., M.O., J.R., X.Y. generated the injected cells and acquired the raw neuronal imaging data of this study. W.Y. helped generate the molecular profiling experiments. M.O. helped optimize the immunolabeling strategy of injected cells. K.C. implemented the auto-tracing platform with assistance from Y.L. K.C. and Z.Y. prepared tracing-related materials, while Y.L. conducted morphological, transcriptomic, and pathological analyses. Other collaborating surgeons provided *ex vivo* tissues and various discussions. Y.L. and H.P. wrote the manuscript with input from all authors.

## Supplementary Figures

**Supplementary Fig. S1.**
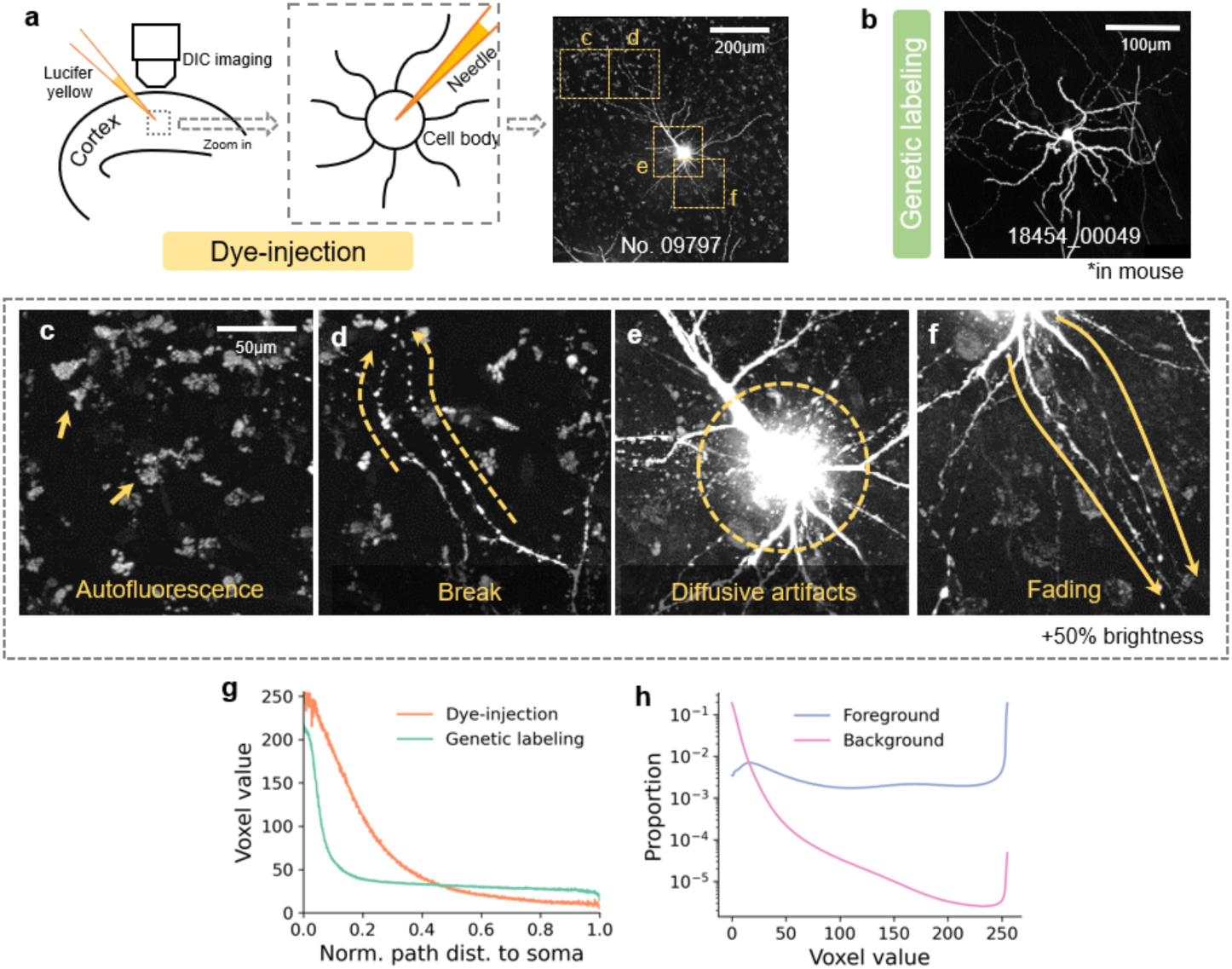
Challenges in human neuron reconstruction. **a**, Schematic of dye injection into a cortical cell body via micropipette. Maximum intensity projection (MIP) of image volume No. 02376 with four regions of interest (yellow boxes) highlighted for subsequent detailed analysis. **b**, MIP of a typical genetically labeled mouse neuron. **c–f**, Common imaging artifacts in human neurons: autofluorescent noise (**c**), signal discontinuities (**d**), perisomatic diffusion halos (**e**), and distal signal attenuation (**f)**, displayed at +50% brightness. **g**, Neurite voxel intensity profiles as a function of normalized distance to soma, comparing dye-injected human neurons (orange) with genetically labeled mouse neurons (green). Voxel values range from 0–255. **h**, Voxel intensity distributions for neurite signal (blue) and background (pink).

**Supplementary Fig. S2.**
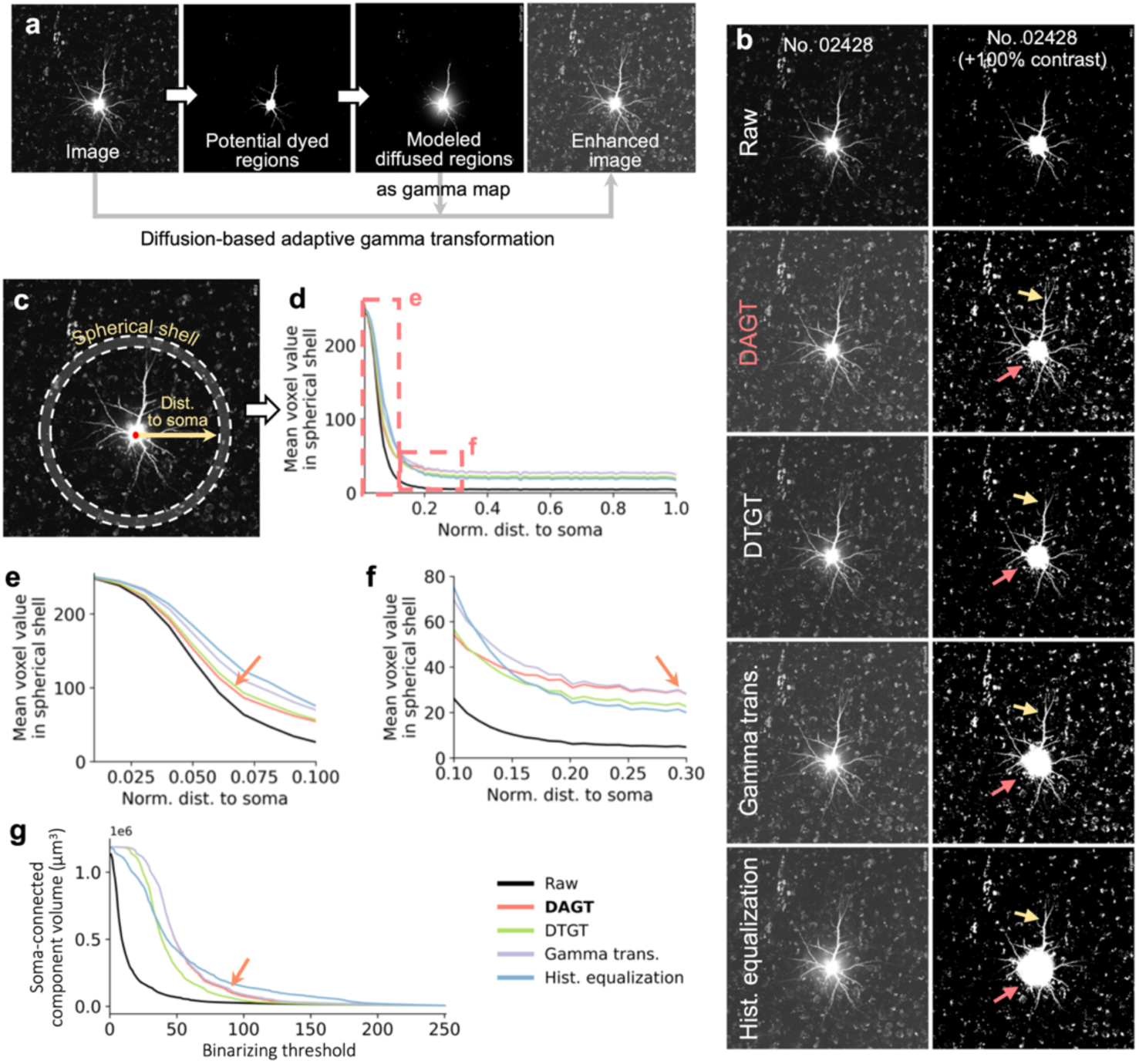
Spatially adaptive ACT-specific enhancement. **a**, Pipeline for diffusion-based adaptive gamma transformation (DAGT) in LetsACT. **b**, Comparison of original and enhanced images using DAGT, derivative-truncated gamma transformation (DTGT), standard gamma transformation, and histogram equalization. Orange arrows indicate high-intensity soma regions; yellow arrows mark weak fibers prone to discontinuities. **c**, Schematic of Sholl-like radial intensity profiling. **d**–**f**, Voxel intensity distributions as a function of normalized distance to soma (**d**), with magnified views of proximal (**e**) and medial regions (**f**). Distance to soma is normalized for each neuron. Orange arrows highlight regions where DAGT performs superiorly. **g**, Volume of soma-containing connected components across binarization thresholds.

**Supplementary Fig. S3.**
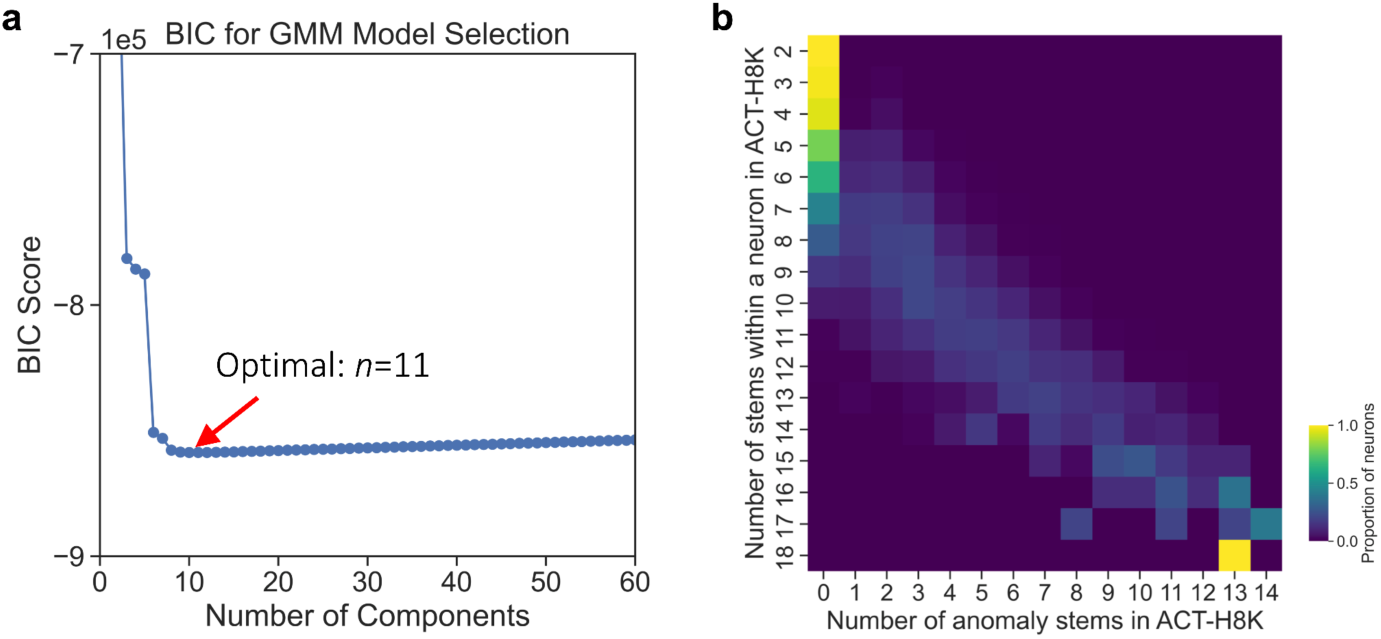
Anomaly stem detection using Gaussian Mixture Modeling of H01. **a**, Bayesian Information Criterion (BIC) values for GMM component selection. The optimal component number (n = 11) is highlighted. **b**, Heatmap showing the relationship between detected anomaly stems and total stem count in initial ACT-H8K-I neurons, with colormap indicating the proportion of neurons in each row.

**Supplementary Fig. S4.**
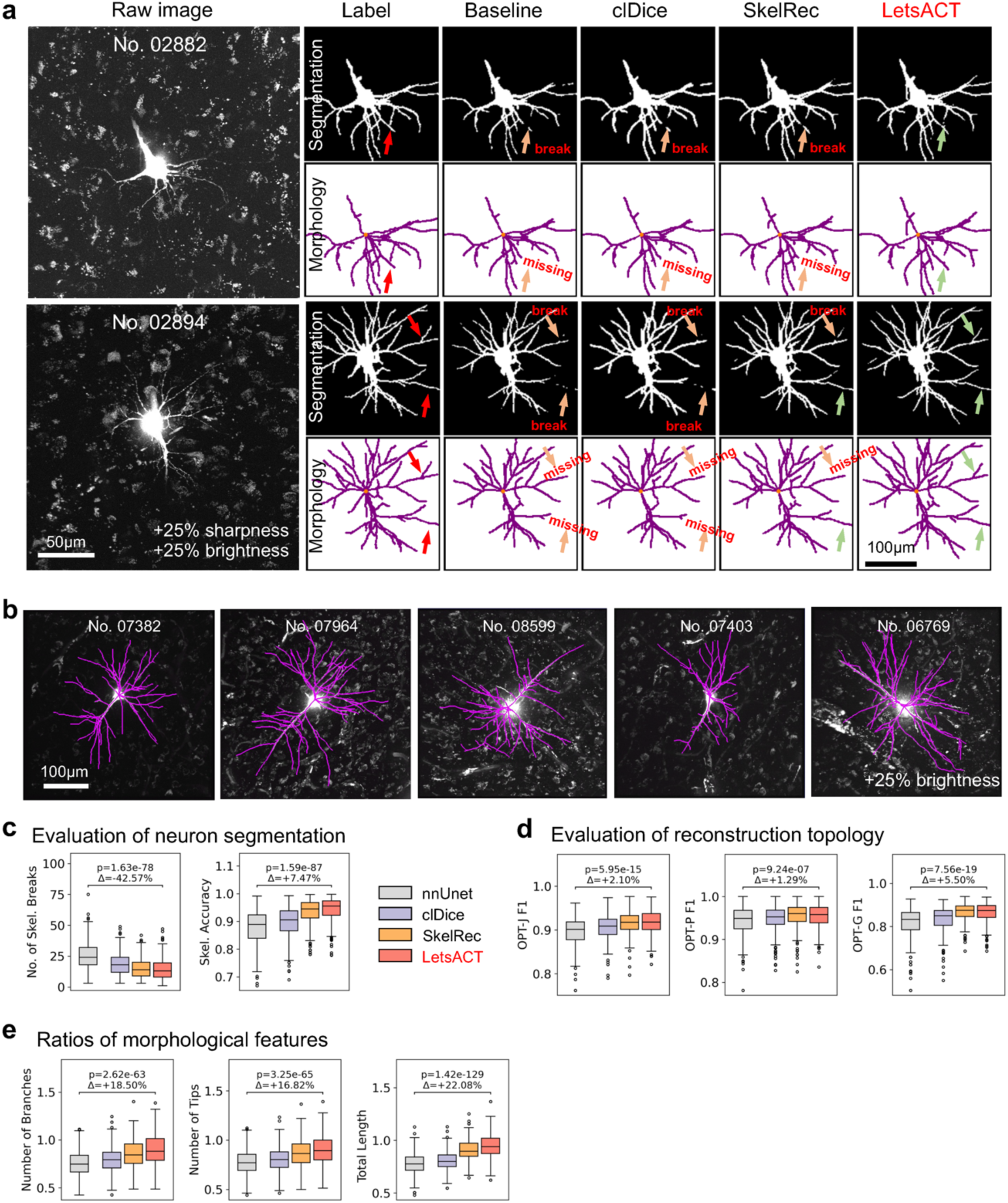
Connectivity-enhancing loss improves segmentation and tracing quality. **a,** Comparison of segmentations and reconstructions across different methods and the manual annotations for two neurons (top and bottom). Neuron IDs are displayed at the top of the raw images. Arrows indicate break-prone fibers (“break”): orange for incorrect segmentation or tracing (“missing”), and green for correct segmentation or tracing. Scale bars: 50 µm for left panels, 100 µm for right panels. **b**, Maximum intensity projection (MIP) of 5 exemplar reconstructions (purple) overlaid onto the images. Scale bar: 100 µm. **c,** Box plots showing segmentation metrics across methods, evaluated by the number of skeleton breaks (left), and skeleton accuracy (right). p: p-value based on two-sided paired t-test, Δ: the improvement of LetsACT compared to the baseline (nnUNet). **d**, Comparison of topological scores (OPT-J, OPT-P, OPT-G) across methods. **e**, Morphological feature ratios versus manual gold-standard annotations.

**Supplementary Fig. S5.**
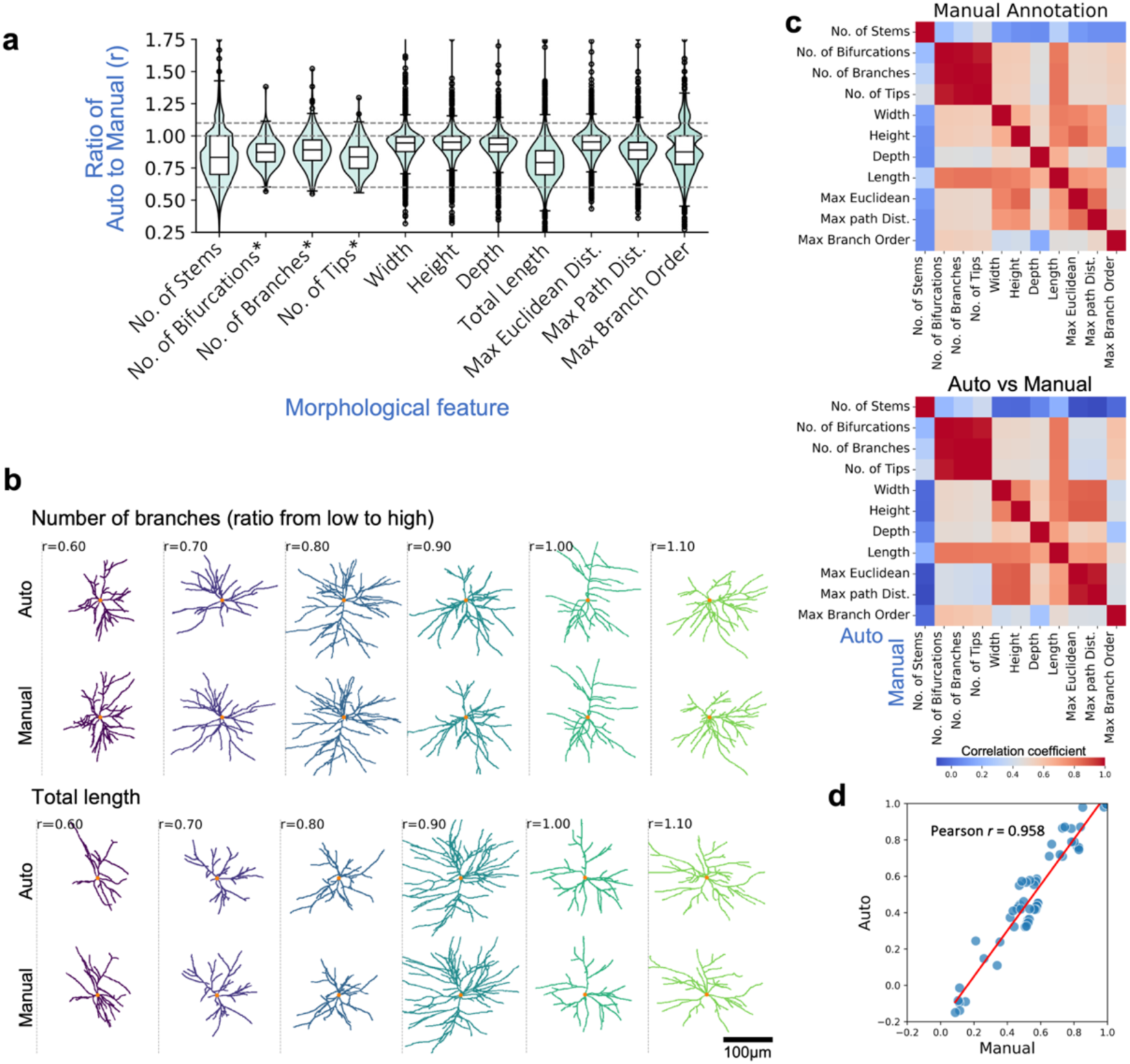
Quality assessment of automated reconstructions. **a**, Violin-and-box plots comparing automated to manual reconstruction ratios across morphological features. Asterisks (*) indicate spatially scaled features, in which each feature value is multiplied by one minus the ratio of its path distance from the soma to the maximum path distance within the neuron. Feature ratios cluster near unity (dashed line, y = 1.0), indicating high concordance between methods. **b**, Visual comparison of manual (top) versus automated (bottom) reconstructions across different accuracy ratios for branch number (top panels) and total branch length (bottom panels). Ratios range from 0.6 to 1.1. Scale bar: 100 μm. **c**, Correlation matrices of morphological features for manual reconstructions (top) and between manual and automated methods (bottom). **d**, Linear correlation between morphological features from automated and manual reconstructions.

**Supplementary Fig. S6.**
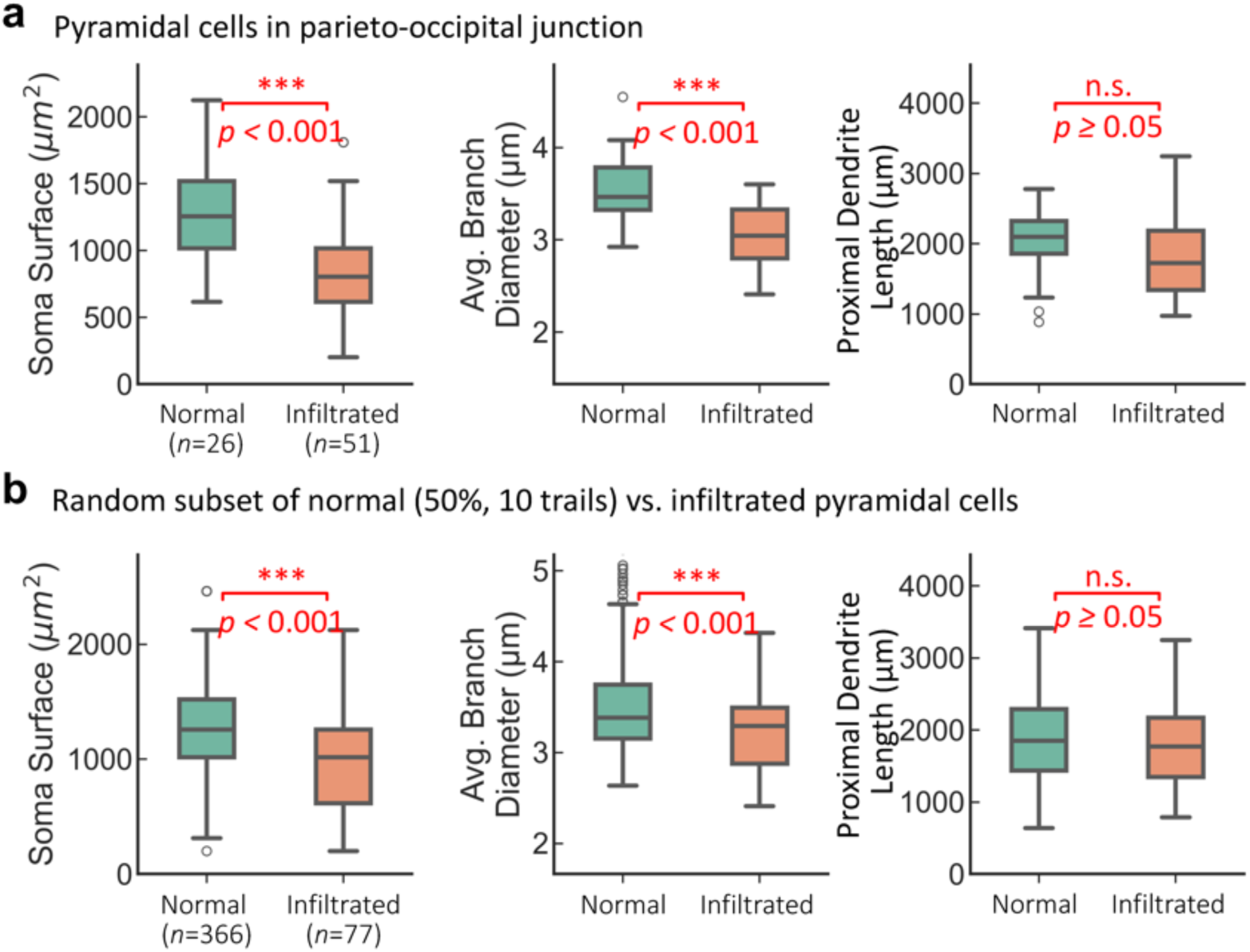
Statistical analyses of pyramidal morphologies from normal and tumor-infiltrated tissues. **a**, Morphological comparisons of pyramidal neurons from parieto-occipital junction: normal tissues (P65, n = 26) versus tumor-infiltrated tissues (P66, n = 51). **b**, Assessment of statistical robustness. Feature statistics were averaged across 10 trials. For each trial, a random 50% subset of normal pyramidal neurons was selected for comparison against the infiltrated neurons (n=77). Statistical significance determined by two-sided Mann-Whitney U test: n.s., not significant; *p < 0.05; **p < 0.01; ***p < 0.001.

**Supplementary Fig. S7.**
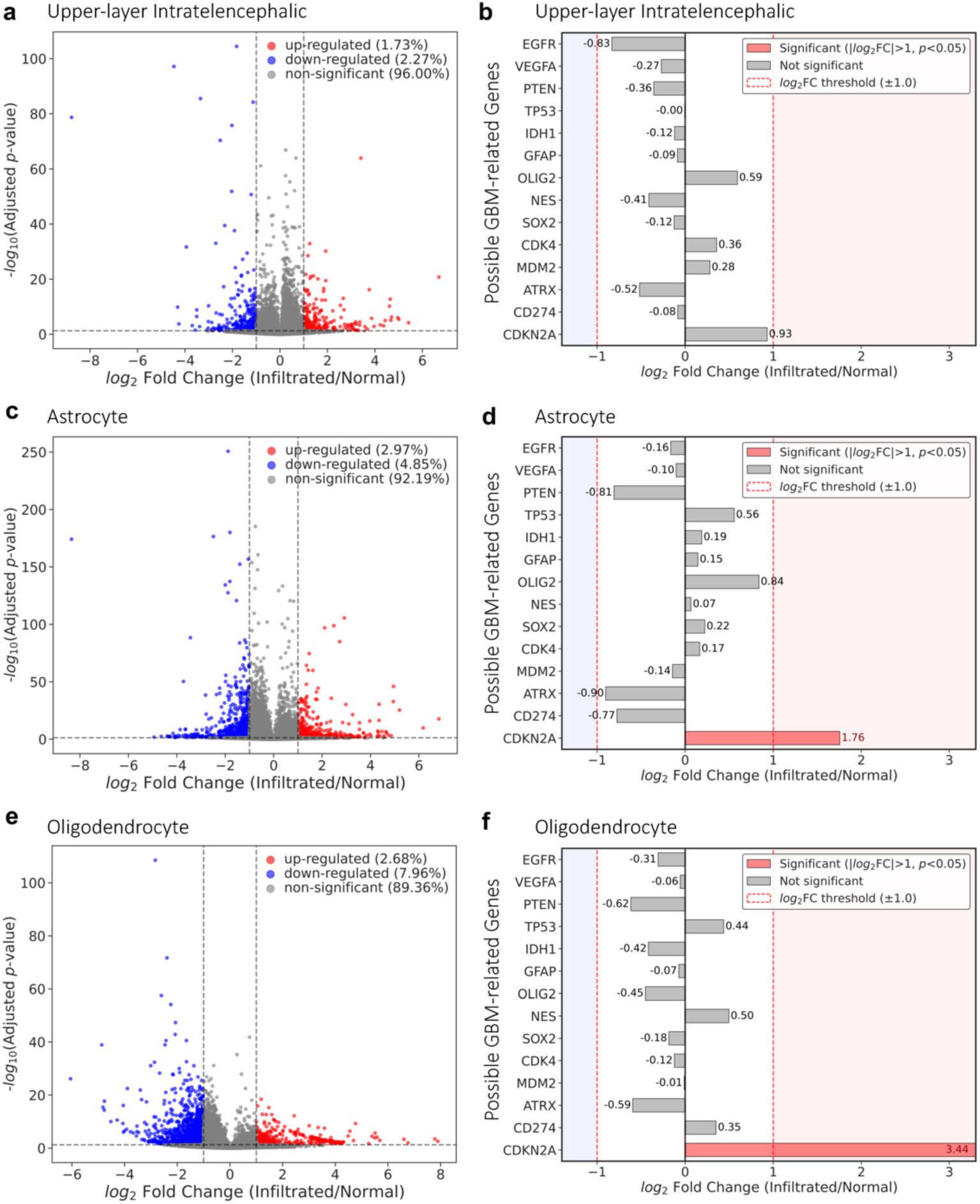
Cell-type specific differentially expressed genes (DEGs) in infiltrated tissues of in-house ACT-10×. **a**, Volcano plot illustrating the global differential gene expression between predicted upper-layer intratelencephalic spots of infiltrated tissue (n=126, P66) and normal controls (n=147, P65) from the junction area of parieto-occipital cortex. Each dot represents a single gene mapped by its magnitude of change (log_2_FC, x-axis) and statistical significance (-log_10_ adjusted p-value, y-axis). Red points indicate significantly up-regulated genes (log_2_FC > 1, p_adj_ < 0.05), and blue points indicate down-regulated genes (log_2_FC < -1, p_adj_ < 0.05). The vertical dashed lines denote the 2-fold change threshold, and the horizontal dashed line marks the significance cutoff (p_adj_ < 0.05). **b**, Differential expression analysis of selected potential GBM-related genes. Bars represent the log_2_FC for each gene. Gray bars indicate genes that did not meet the criteria in this cohort (|log_2_FC| > 1, p_adj_ < 0.05), despite varying levels of statistical significance (asterisks denote significance: *p < 0.05, **p < 0.01, ***p < 0.001, n.s. = not significant). **c-f**, Similar volcano plots and bar plots for astrocyte spots (**c**, **d**; infiltrated: n=823; normal: n=451) and oligodendroctye (**e**, **f**; infiltrated: n=854; normal: n=161).

**Supplementary Fig. S8.**
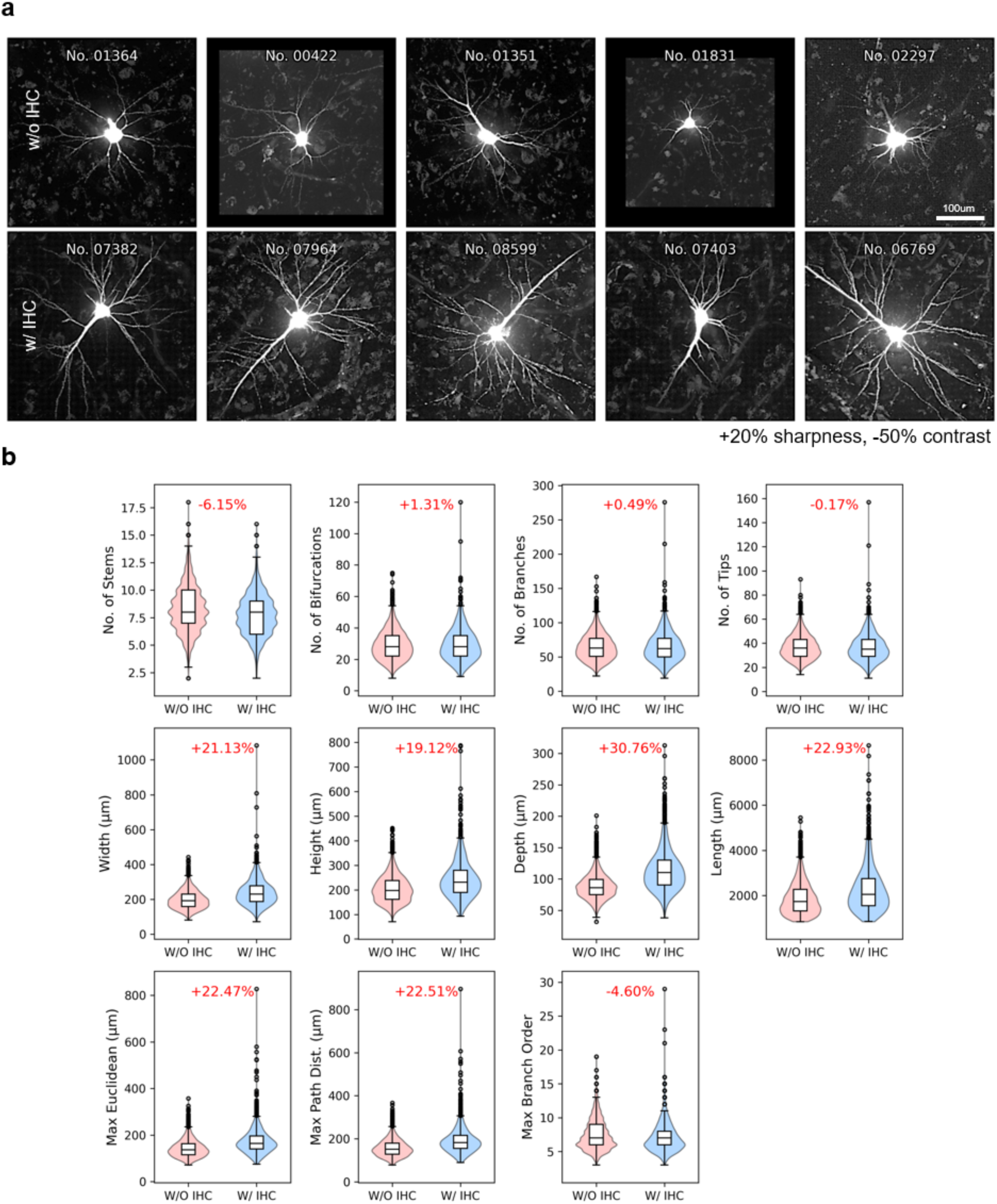
Comparison of neurons with and without secondary immunohistochemical staining. **a**, Representative neuronal MIP images showing increased brightness and neurites using IHC staining. **b**, Morphological feature distributions of reconstructions from non-IHC-stained versus IHC-stained neurons, with percent increase shown in red. While branching patterns remains largely unchanged (number of bifurcations, branches, and tips), branch lengths increase (total dendrite length, maximum Euclidean distance, maximum path distance). This effect is most pronounced in distal branches, resulting in larger overall reconstructions (width, height, depth).

**Supplementary Fig. S9.**
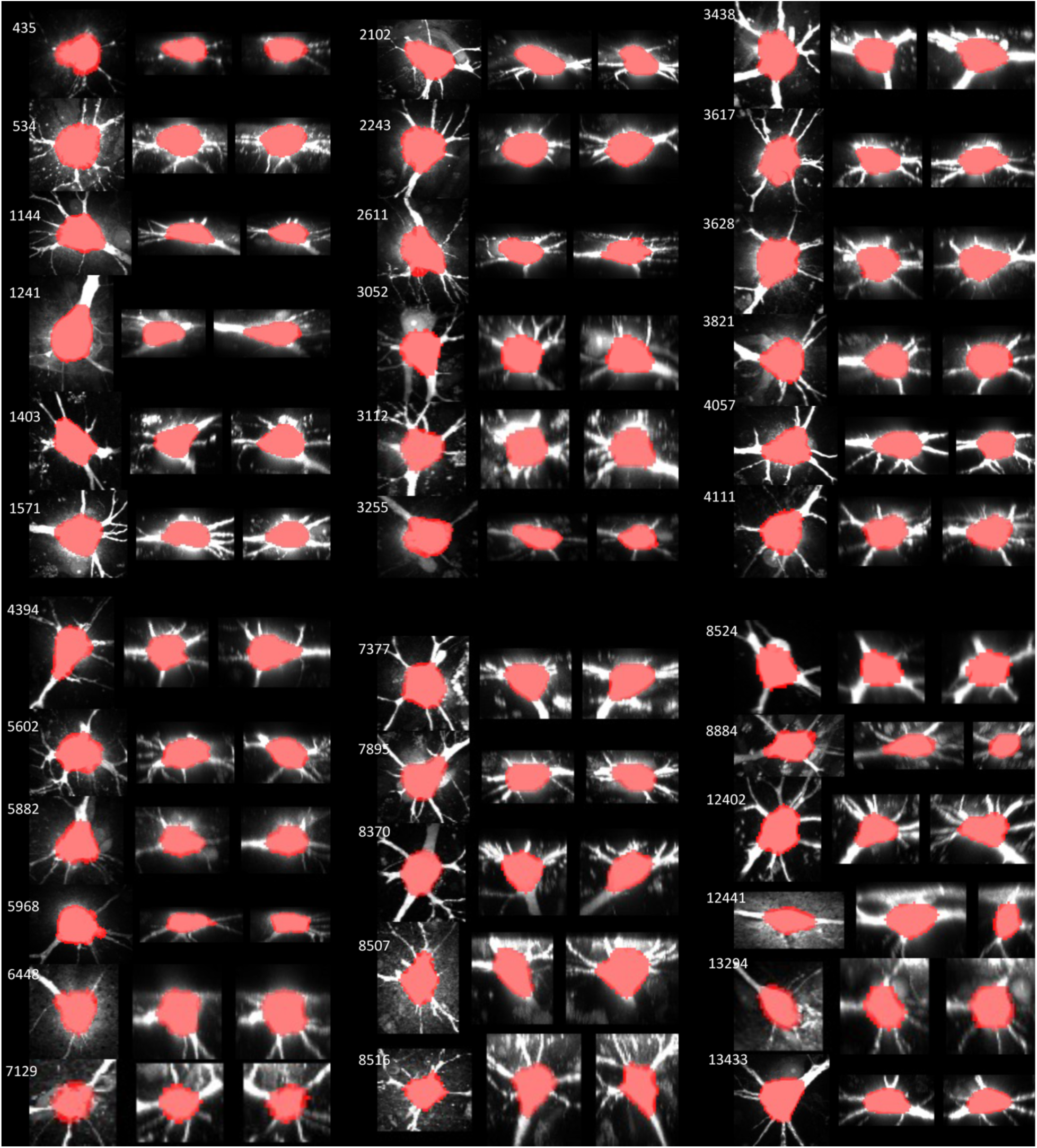
Random examples of soma segmentation masks. Cell IDs are indicated at the top left of each panel set. Each neuron is visualized through three orthogonal views: the xy plane (left), xz plane (middle), and yz plane (right). The white signal represents the maximum intensity projection (MIP) of the soma-centered region, while the orange overlay delineates the corresponding segmentation mask.

## Supplementary Tables

**Supplementary Table S1. | Metadata for surgical tissue samples.**

**Supplementary Table S2. | Metadata for reconstructed neurons.**

**Supplementary Table S3. | Annotated cell types.** Cell type annotations for ACT-H8K neurons, including cell identifier (“name”), cell type classification (“CLS2”), and number of annotators (“num_annotator”). Cell type codes: 0, pyramidal; 1, nonpyramidal; 3, conflicting annotations (uncertainty between annotators); –, undetermined.

